# Orally Delivered Milk-Derived Nanovesicles Loaded with Connexin 43 Peptides for Targeted Cardiac Ischemia-Reperfusion Therapy

**DOI:** 10.1101/2025.01.01.630994

**Authors:** Spencer R. Marsh, Md Ruhul Amin, Stefano Toldo, Claire Beard, Alan B. Dogan, Eleonora Mezzaroma, Ethan Andres, Randy F. Stout, Mark S. Bannon, Laura Beth Payne, Antonio Abbate, Yassine Sassi, Rachel A. Letteri, Robert G. Gourdie

## Abstract

Extracellular vesicles have emerged as promising nanocarriers for targeted drug delivery, but their therapeutic potential is limited by challenges related to administration route, loading, targeted delivery and production at scale. Here, we report an innovative approach for targeted delivery of therapeutic peptides to injured tissues using milk-derived small extracellular vesicles (mEVs) as an abundant, safe, orally administrable nanoplatform. We demonstrate that a sub-population of mEVs naturally contain Connexin 43 (Cx43) and its Carboxyl-Terminal (CT) polypeptides, which have been shown to play crucial roles in wound healing and tissue repair. Leveraging this intrinsic property, we developed an esterification method to efficiently and uniformly load mEVs with enhanced levels of an exogenous Cx43 CT peptide (αCT11 - RPRPDDLEI), as assessed by flow cytometry-based vesicle quantification and mass spectrometry. These engineered mEVs exhibited remarkable injury targeting capabilities, with > 30-fold increases in uptake by injured cells compared to non-wounded cells in vitro and preferential accumulation in wounded tissues in vivo. Notably, αCT11-loaded mEVs orally administered after myocardial infarction reduced infarct size by >60% and preserved heart function in a mouse model of ischemia-reperfusion injury. This study represents a significant advance in nanomedicine, demonstrating the utilization of naturally occurring milk-derived extracellular vesicles as an oral delivery system for therapeutic peptides, achieving unprecedented targeting efficiency and efficacy in the treatment of myocardial ischemia-reperfusion injury.

## Introduction

Small extracellular vesicles (EVs) are lipidic nanovesicles, 50-200 nm in diameter, secreted by nearly all eukaryotic cells ^1,2^. Among EVs, exosomes are distinguished by their biogenesis through the Endosomal Sorting Complex Required for Transport (ESCRT) mechanism and specific molecular markers such as CD9, CD81, Syntenin, and TSG101^3^. The composition of EVs can vary significantly, reflecting cellular origin and physiological state, which influences their roles in intercellular communication and disease processes ^4–6^.

EVs have emerged as promising nanocarriers for drug delivery, but their therapeutic potential is limited by challenges in administration route, drug loading, targeted delivery and limitations on the ability to produce EVs at a scale required for pharmaceutical translation ^7,8^. Milk-derived EVs (mEVs) offer a unique solution to these challenges due to their natural abundance, safety profile, stability, ability to be orally administered and high bioavailability once ingested ^9–12^. Additionally, mEVs invoke minimal immune responses, demonstrate reduced clearance by first-pass metabolism, and an ability to cross biological barriers like the blood-brain barrier – all features that enhance their therapeutic potential ^13–15^.,

Connexin 43 (Cx43), a gap junction protein widely expressed in mammalian tissues, is present in EVs and plays crucial roles in wound healing and tissue repair ^16,17^. The carboxyl-terminal (CT) domain of Cx43, including its alternately translated isoform GJA1-20k ^18,19^, has shown significant therapeutic potential in cardiac injury responses and disease ^20,21^. Synthetic peptides containing the Cx43-terminus (CT) sequence RPRPDDLEI (αCT11), incorporated in GJA1-20k, have also demonstrated efficacy in reducing ischemic injury in ex vivo models ^22^ and enhancing wound healing, including in clinical tests as an external topical treatment for diabetic foot and venous leg ulcers ^23–25^. However, internal administration of αCT11 in vivo is limited by the fact that this short linear peptide breaks down within 10 minutes in blood or serum ^26^, necessitating an approach to sustaining bioactivity long enough to mediate clinically meaningful effects.

In this study, we present a new approach to utilizing milk-derived extracellular vesicles (mEVs) as a natural, orally administrable nanoplatform for targeted delivery of therapeutic peptides to injured tissues. We demonstrate that a subset of mEVs inherently contains full-length Connexin 43 (Cx43) and C-terminal (CT) polypeptides. Exploiting this intrinsic property, we developed a novel esterification method to efficiently load an exogenous Cx43 CT peptide (αCT11 - RPRPDDLEI) into nearly the entire isolated mEV population. This peptide has previously shown anti-arrhythmic and cardioprotective effects^22,27,28^. Our findings demonstrate that mEVs exhibit natural injury-targeting properties, with increased uptake in both cardiac ischemic injuries and skin wounds in mice. Leveraging this platform, we demonstrate, for the first time, the successful oral administration of a therapeutic peptide that exhibits significant cardioprotective efficacy when administered post-myocardial infarction in a murine model of cardiac ischemia-reperfusion injury. Our drug delivery technology could facilitate non-invasive, targeted delivery of various therapeutic peptides and biologics that have shown preclinical promise but have faced challenges in clinical translation.

## Results

### Characterization of Small Extracellular Vesicles (EVs) Isolated from Bovine Milk

Figure 1 summarizes the method used to isolate, purify and validate EVs from bovine milk (mEVs). The approach shown revises and updates our previously published method ^10^, which uses EDTA to free mEVs from milk protein complexes, including casein micelles^10^. Among other innovations, the method uses Tangential Flow Filtration (TFF at <500 kDa) to harvest disassociated mEVs, thereby eliminating the ultracentrifugation (UC) step commonly utilized in most extracellular vesicle (EV) isolation protocols (Fig. 1A). TFF followed by Sepharose column (SEC) fractionation leads to improvements in mEV yield, ultrastructural morphology and intactness over UC-based methods, enabling production of mEVs at scale.

**Figure 1:**
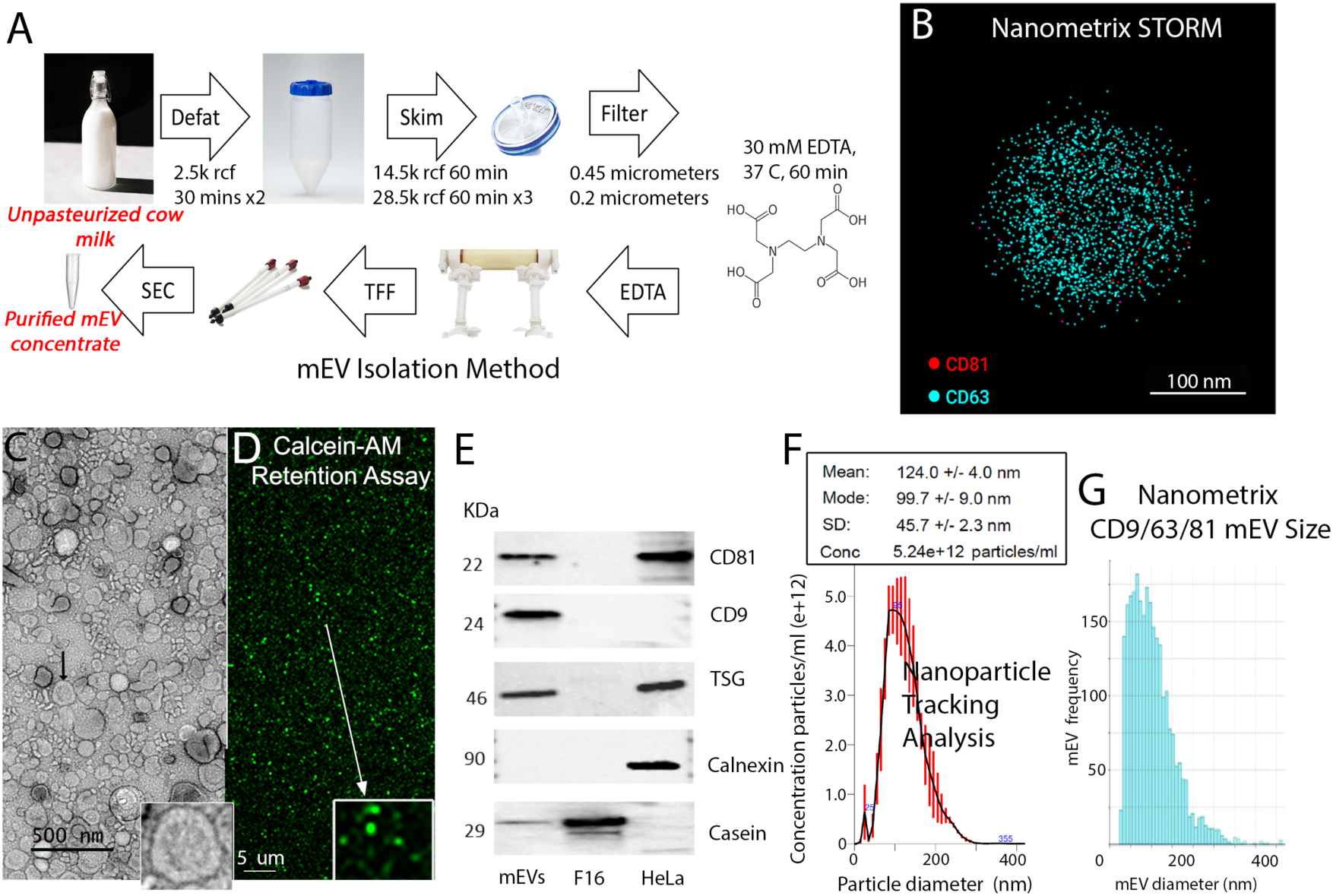
Milk-derived small extracellular vesicle (mEV) characterization. A) Schematic of mEV isolation protocol showing the steps of the procedure. Fractions referred to are those collected during Sepharose Column (SEC) fractionation. Fractions 7 through 9 (F7-F9) show highest levels of mEV enrichment and samples used in this study were pooled from F7-F9. Fractions >14 show enrichment for contaminating milk proteins such as casein, with low or no evidence of mEVs, and are typically discarded. B) Single Molecule Localization Microscopy (SMLM) image of an isolated mEV isolated by the protocol in (A) tagged for CD81 (red) and CD63 (teal). C) Negative stain transmission electron microscopy (TEM) indicating dense accumulations of mEVs generated by the isolation protocol. D) Calcein AM loading assay illustrating membrane intactness of isolated mEVs E) Western blotting of mEV isolates for CD81, CD9, TSG-101, and negativity and markedly reduced expression of Calnexin and Casein, respectively. Lysates from HeLa cells are included as a control. Sepharose fraction 16 (F16) shows elevated casein, but no evidence of EV markers. F) Nanoparticle Tracking Analysis (NTA) of an mEV isolate indicating >5×10^12^ mEVs per mL, with average nanoparticle size of 125.0 nm, consistent with small EVs. G) SMLM-generated size distribution of CD9/CD81/CD63-positive mEVs, showing a similar histogram morphology to NTA results in panel F.

Quality control checks routinely carried out after each mEV isolation include a range of analytical methods –summarized in Figure 1, including Single Molecule Localization Microscopy (SMLM) imaging (Fig. 1B, G), Transmission Electron Microscopy (TEM, Fig. 1C), Calcein AM dye uptake and retention (Fig. 1D), western blotting (Fig. 1E), and Nanoparticle Tracking Analysis (NTA, Fig. 1F).

TEM was used to confirm dense concentrations of mEVs with undisrupted vesicular ultrastructural morphology (Fig. 1C). Esterified Calcein AM (Fig. 1D) was used to assess the patency (i.e., non-leakiness) of mEV isolates. Upon de-esterification, Calcein is retained within the lumen unless the integrity of the mEV membrane is compromised and leads to leakage. Figure 1D shows mEVs maintaining punctate Calcein signals 4-hours after labeling, indicating stable retention of the dye. NTA was used to assay mEV size, counts and histogram shape. Figure 1F shows typical NTA data, with mEVs of mean diameter of 124 nm; a symmetrical, single-peaked size distribution; and nanoparticle counts in excess of 5 ×10^12^/mL.

Figure 1E shows representative western blots for EV markers including CD9, CD81 and TSG-101, together with confirmation of absence of Calnexin, a cell membrane marker found in large extracellular vesicles (apoptotic bodies), but not in small extracellular vesicles ^29^. Casein levels in the pooled EV fraction were also probed by western blotting to confirm reduction relative to later non-EV containing SEC fractions (Fraction 16, F-16) as shown in Figure 1E.

To further characterize mEV isolate morphology and composition, SMLM imaging was undertaken in collaboration with Nanometrix Inc. SMLM enables detection of individual fluorescent molecule emission events, and can achieve <40 nm spatial resolution, providing super resolution images of sub-diffraction-sized objects (i.e., <200 nm), such as small EVs. Figure 1B shows a SMLM image of a milk EV co-labeled for CD63 (teal) and CD81 (red). Histograms of nanoparticles co-labeled for the EV markers CD63, CD9 and CD81 quantified by SMLM are comparable in morphology, size distribution and mean diameter to those obtained by NTA (Fig. 1G). Additional quality tests carried out to rigorously check the pooled mEV fraction after each isolation include Nanodrop assays of protein concentration (expected ∼1.0 mg/mL +/-0.1 mg/mL) and nanoparticle zeta potential (expected ∼-7.0mV +/-1.0 mV).

### Connexin43 Polypeptides are Present in Milk EV Sub-Populations

Western blotting for gap junctional Cx43 was undertaken on mEV isolates - pooled following SEC and characterized as described in the preceding section. Three antibodies against distinct epitopes in the cytoplasmic loop (CL) and CT of Cx43 were used (Figure 2A). A band at ∼40-44 kDa was detected in mEV isolates by the Cx43 CL antibody, consistent with full-length Cx43 ^30^. Similarly, the two Cx43 CT antibodies labeled bands at 40-44 kDa, with multiple bands in this size range consistent with the tendency of Cx43 antibodies to detect multiple Cx43 phospho-isoforms ^17^. However, the Cx43 CT antibodies also detected bands below 40 kDa, most prominently in the 14 to 20 kDa range (Figure 2A). In control lysates from HeLa cells that exogenously express Cx43 (HeLa-Cx43), blotted with the Cx43 CT antibody, prominent bands at 40-44 kDa and 20-22 kDa were observed. This pattern is consistent with the presence of full-length Cx43 (∼43 kDa) and a 20 kDa Cx43 (GJA1-20k) isoform generated by alternative translation of Cx43 mRNAs ^18^. To validate the presence of Cx43 polypeptides, we also performed immunoprecipitation of Cx43 with a CT antibody against amino acids (AAs) 363 to 382 and probed the immunoprecipitated bands with a second CT antibody against AAs 333 to 346 (Fig. 2B). This resulted in detection of strong bands consistent with full-length Cx43 (∼43 kDa) and a 20 kDa Cx43 in mEVs, as well as fainter bands in the 14 to 16 kDa range - see long exposure blot of the IP in Figure 2B.

**Figure 2:**
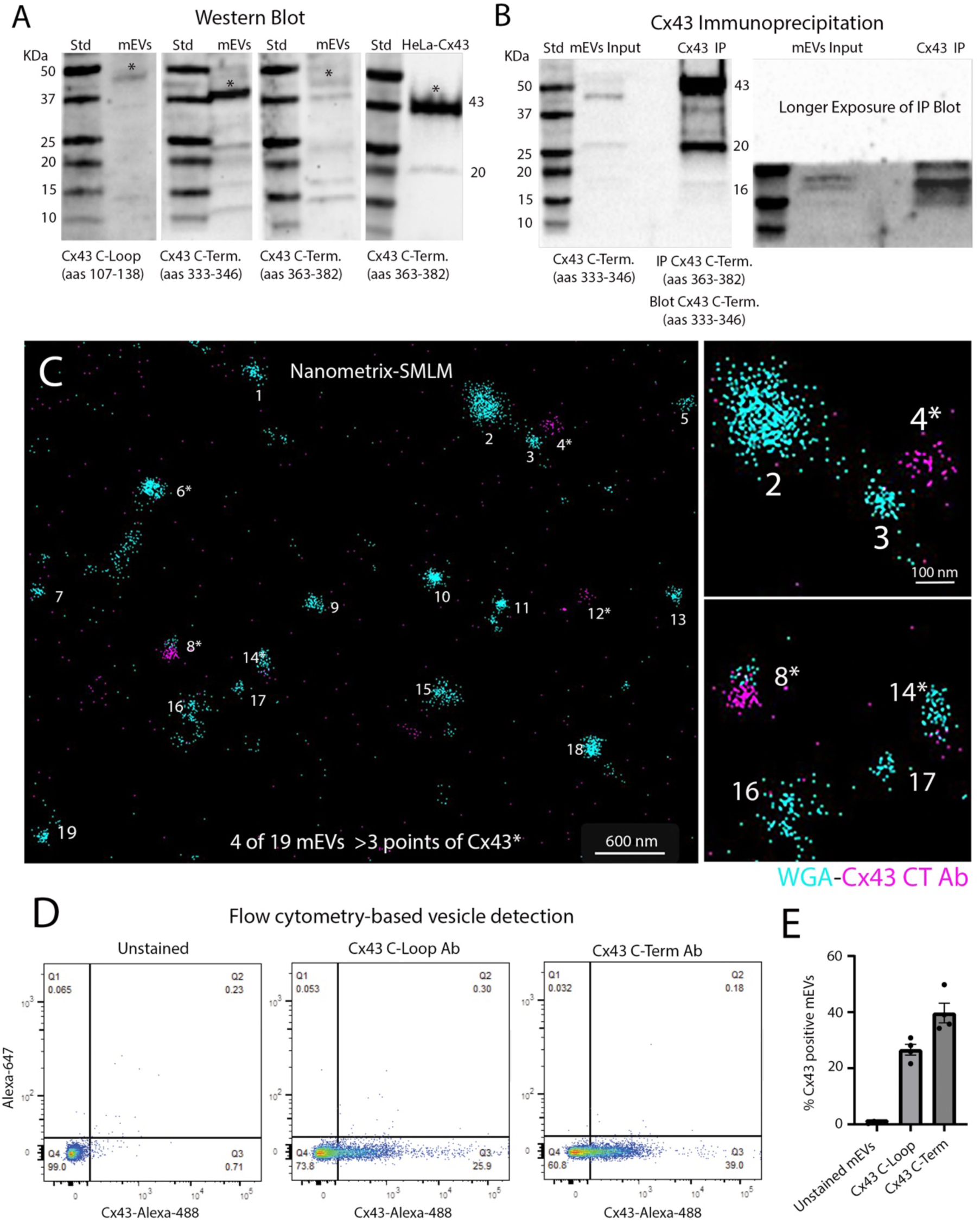
A sub-population of mEVs contain Cx43 polypeptides. A) Western blots for Cx43 in mEVs and control lysates in HeLa cells heterologously expressing Cx43. The 3 antibodies used are against a Cx43 cytoplasmic loop (CL) peptide epitope (AAs 107-138) and two different epitopes on the CT of Cx43 (AAs 333-346 and 363-382). Single asterisk indicates bands at ∼43 kDa detected by each antibody. Note the presence of multiple bands below 25 kDa detected by both Cx43 CT antibodies in mEVs. Full-length 43 kDa and shorter 20 kDa Cx43 CT isoforms are also present in control HeLa-Cx43 cells. B) Immunoprecipitation (IP) of Cx43 in mEVs with the CT antibody against amino acids 363 to 382 and blotted with the antibody against Cx43 CT amino acids 333 to 346. The lower part of the IP Blot was exposed for a longer time-period to highlight lower molecular mass bands below 25 kDa present in the mEVs. C) SMLM super-resolution imaging of mEVs detected by a Cx43 CT antibody (red), together with wheat germ agglutinin (WGA-teal) co-labeling. D) Flow cytometry-based vesicle detection of mEVs using antibodies against the Cx43 C-Loop (AAs 107-138, right hand plot in D) and C-Term (AAs 363-382, left hand plot in D) directly conjugated to Alexa-488 fluorophores. Unstained mEV controls were used to set the fluorescent gates. E) Quantification of flow cytometry-based vesicle detection of mEVs positive for Cx43 polypeptides, as detected by the C-Loop and C-Term Cx43 antibodies. The Cx43-positive sub-population appears to comprise 20-40 % of the pool of total mEVs. N = 3 experimental replicates for each group, error bars= SEM.

SMLM super-resolution imaging was undertaken on mEVs using a Cx43 CT antibody (AAs 363-382), together with wheat germ agglutinin (WGA) co-labeling of mEVs (Fig. 2C). Some (∼20 % of WGA+ mEVs), but not all, mEVs showed evidence of Cx43 signals. To further investigate the Cx43-positive sub-population vesicle we undertook flow cytometry-based vesicle detection of mEVs ^31^. Detection in flow cytometry was performed using the Cx43 CT (AAs 363-382) and Cx43 CL antibodies (AAs 107-138) directly conjugated to Alexa-488 fluorophores (Figure 2D). A summary of the gating steps for Cx43 flow cytometry-based vesicle detection is shown in Supplemental Figure 1. Unstained mEVs showed minimal background fluorescence and were used to set the fluorescent gates (Figure 2D, LH panel). mEVs single-stained for either Cx43 CL or Cx43 CT antibody displayed 20 to 40% positivity (Figure 2D and 2E). Thus, both SMLM and flow cytometry-based vesicle detection, indicated the presence of a sub-population within the broader population of mEVs that was positive for Cx43 polypeptides.

**Supplemental Figure 1:**
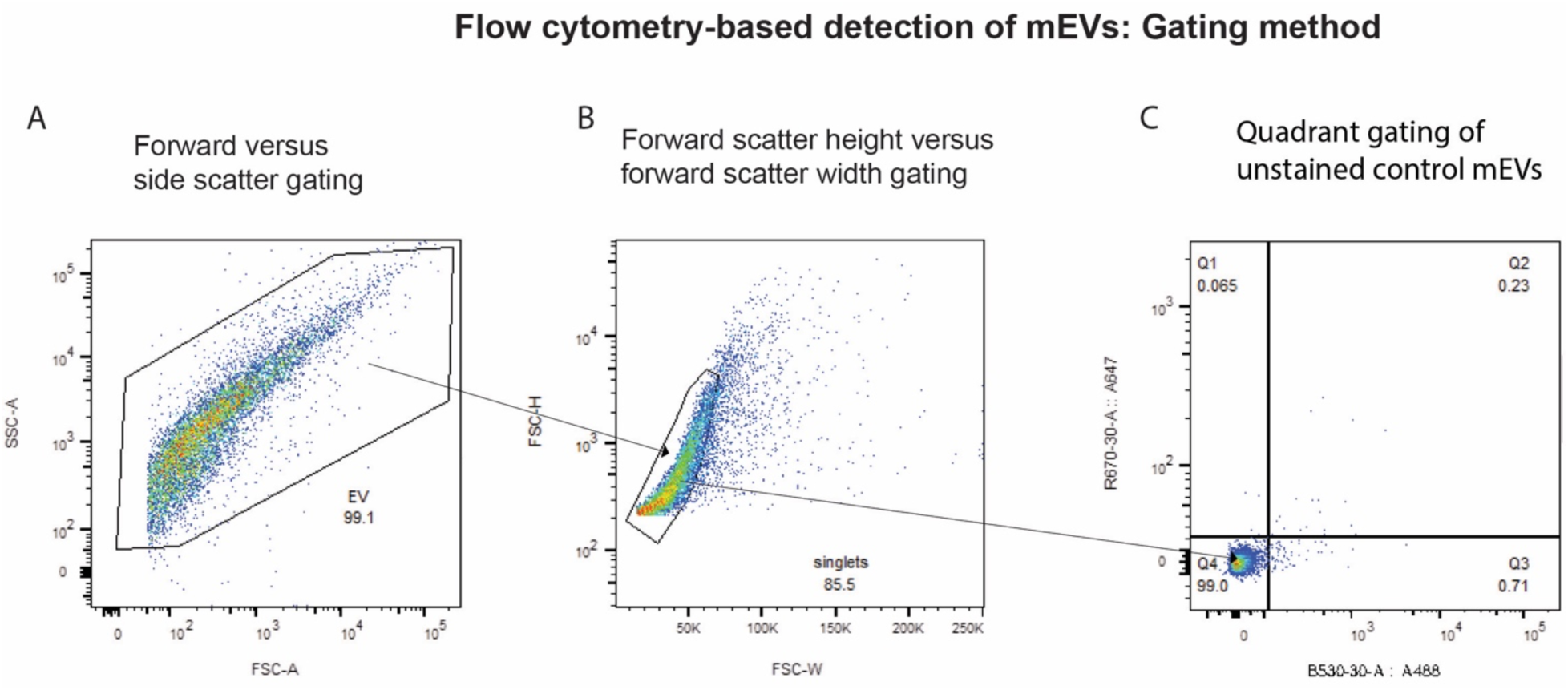
Gating method of flow cytometry-based detection of unstained mEVs: Samples were collected on a Becton Dickinson Fluorescence-Activated Cell Sorting Fusion Flow Cytometer. A) Unstained EVs were displayed on a Forward Scatter Area (FSC-A) and Side Scatter Area (SSC-A) dot plot using log scale and a gate was drawn around the EV events. B) A subsequent plot of Forward Scatter Height (FSC-H – log scale) vs Forward Scatter Width (FSC-W-linear scale) was displayed and gated to remove aggregate events. C) Acquisition was set to record 10,000 singlet events per sample and unstained mEVs were used to set the quadrant gating for potential Alexa-647 vs Alexa-488 signals.

### Milk EVs Prompt Faster Healing of Wounded Monolayers of Dermal Fibroblasts

Cx43 CT polypeptides, such as the endogenous GJA1-20K, as well as the synthetic Cx43 CT fragment αCT1, have been found to modulate injury and wound healing responses ^25,32^. Given that multiple Cx43 isoforms of 20 kDa or below were detected (i.e. Figure 2A, B), we sought to determine whether mEVs possessed bioactivity in vitro in scratch-wound assays, resembling those associated with Cx43 CT polypeptides^33,34^.

Human dermal fibroblasts (HDefs) were cultured in monolayers to confluency then mechanically wounded with a pipette tip, rinsed, and provided with fresh media with either vehicle control (HEPES buffer), 20 μg/mL of mEVs, or 100 μM αCT1 as a positive control (Fig. 3A, B). An image of the initial scratch area (Post-Wound) for each group is shown in the top row of Figure 3A, while final scratch area (T=6 hours post-treatment) for each group is shown in the lower row. Migration index was then calculated for each group – ((Initial Wound Area-Final Wound Area)/Initial Wound Area) x 100 – normalized to the vehicle control group), as we have previously reported ^33,34^. Cells treated with mEVs showed qualitatively and statistically significant (p<0.001) increases in the rate of closure of scratch wounds relative to vehicle controls (Figure 3B), reaching a level matching 40% that of αCT1, which was also significantly elevated above control (p<0.0001).

**Figure 3:**
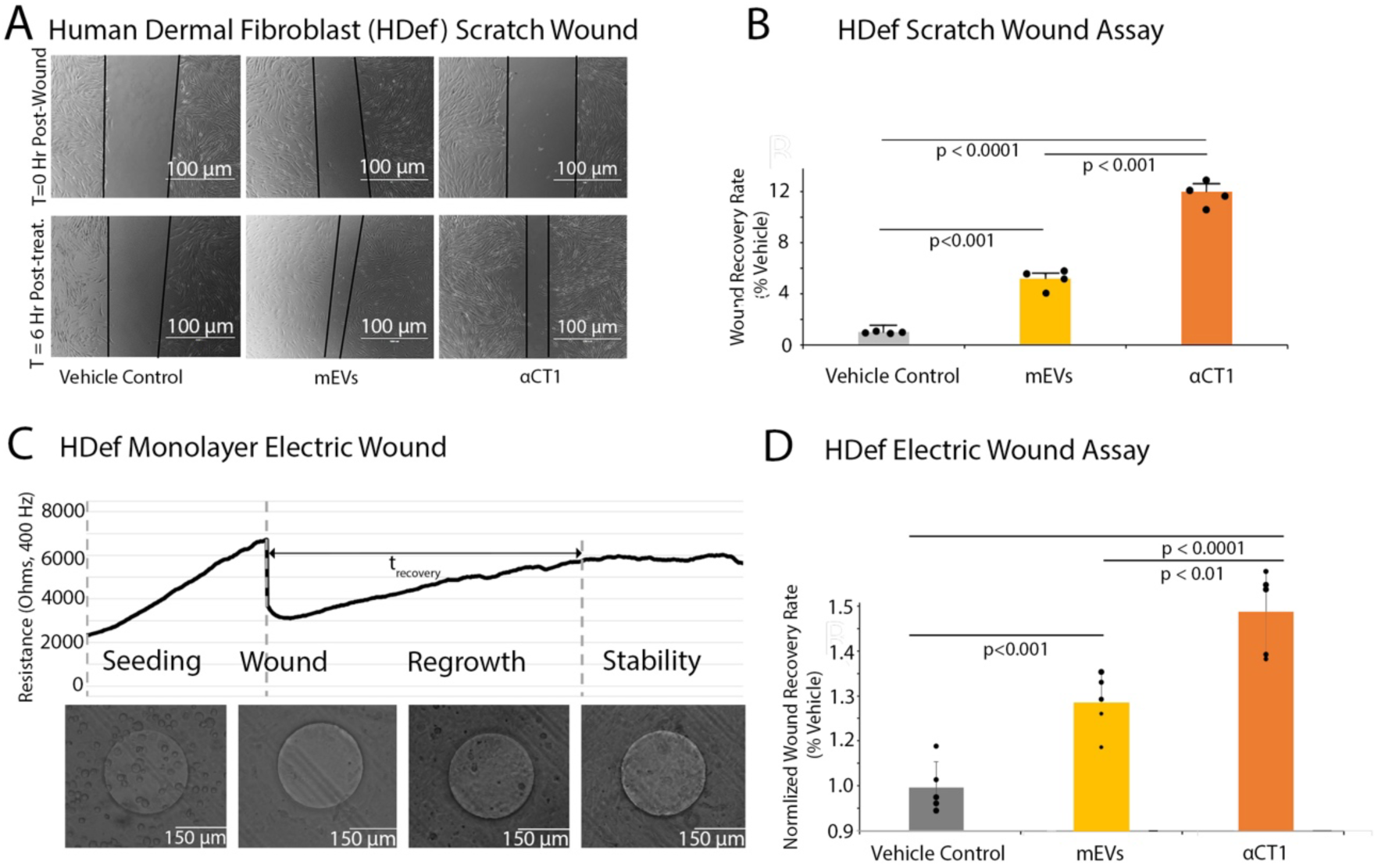
mEV effects on wound healing in human dermal fibroblasts (HDef). A) Scratch wound images from t=0 hours (post-wound) and t=6 hours (post-treatment), indicating level of regrowth of HDef monolayers in mEV-treated cultures relative to vehicle controls. Vehicle control is addition of HEPES buffer only to cultures; mEV treatment is addition of mEVs at 20 µg/mL in HEPES buffer; and an αCT1 peptide positive control is addition to culture media of the (non-EV encapsulated) peptide in HEPES buffer at 100 μM. B) Quantification of healing rates from HDef scratch wound assays, indicating significant acceleration of wound healing in mEV- and 100 μM αCT1-treated monolayers over the vehicle control group. N= 4 experimental replicates for each group, Error = SEM. C) Representative ECIS resistance trace of HDef monolayers on ECIS electrodes during an electric wound experiment: 1) the “Seeding” phase, where fibroblasts form a monolayer over the electrode; 2) the “Wound” phase, where an elevation in electrical current clears cells from a circular 200 μM diameter region over electrode; 3) the “Regrowth” phase, where cells reform a monolayer over the electrode; and the “Stability” phase, where cells reach confluence. Resistance reflects the proportional coverage of the ECIS electrode over the 6–8-hour time course of the experiment. D) Quantification of healing rates of HDefs in ECIS electric wound assays for vehicle controls, mEVs, and 100 μM αCT1 treatments as in (B) – Normalized Wound Recovery Rate is a function of treatment Time to Recovery (T_recovery_), as defined on the ECIS graph in (C) relative to the vehicle control mean. As in (B), mEV- and αCT1-treated HDef cells show significant acceleration of regrowth of cells over the electrode relative to the vehicle control group. N = 5 experimental replicates for each group, Error = SEM.

To determine whether these effects persisted in a complementary injury scenario in vitro, we used an automated protocol based on Electric Cell Impedance Sensing (ECIS), which tracks migration of cells into a circular 200 μm diameter patch cleared by electrical wounding of the monolayer (Fig. 3C). The pattern observed for electrical wounding of HDef monolayers using ECIS was similar to that of mechanical injury. Over the first 2-4 hours, both mEV- and αCT1-treated cultures showed accelerated migration into the area of electrically cleared cells relative to the vehicle control. This trend culminated in both αCT1 and mEV-treated cells migrating into the cleared region of electroporated cells significantly faster over the time course, relative to the vehicle control (p<0.001) (Fig. 3D). As was the case with the scratch-wounding assay, treatment with αCT1 had the largest migration-promoting effect on closure of ECIS wounds in HDef monolayers, promoting a statistically (p<0.01) increased rate of closure over milk EVs (Fig. 3D).

### Cellular Cx43 Expression Affects mEV Acceleration of Wound Healing in Vitro

Next, we sought to determine whether Cx43 expression affected the wound healing response in mEV-treated cells, as observed in Figure 3. We repeated scratch wounding on monolayers of Madin Darby Canine Kidney (MDCK) cells lacking endogenous Cx43, as well as on MDCK cells that had been stably transfected to express wild-type rat Cx43 (Fig. 4)^35,36^. Immunolabeling and Western blotting of MDCK Cx43- and Cx43+ cells confirmed the absence and high expression of Cx43 in these lines, respectively (Fig. 4A, B). Cx43+ and Cx43-MDCK cultures were grown to confluency and mechanically wounded (Fig. 4C, D), with initial scratch area (post-wound) for each group shown in the upper panels, and scratch area 6 hours post-treatment is shown in the lower row of panels. Bar graphs indicating relative migration index for experimental groups are shown in Figure 4E (Cx43-cells) and Figure 4F (Cx43+ cells). Migration index for Cx43- and Cx43+ MDCKs was similar and low (∼1) in the presence of vehicle control solution. Addition of milk EVs to scratched Cx43-MDCK monolayers resulted in a significant (p<0.001), albeit modest increase in migration – ∼2-fold higher than vehicle control (Fig. 4E). By contrast, addition of milk EVs to scratched Cx43+ MDCK monolayers resulted in an ∼10-fold increase in migration relative to control non-EV-treated Cx43+ MDCKs (p<0.001) or Cx43-MDCKs (Fig. 4F), suggesting that Cx43 expression increased cell-migration response to mEVs.

**Figure 4:**
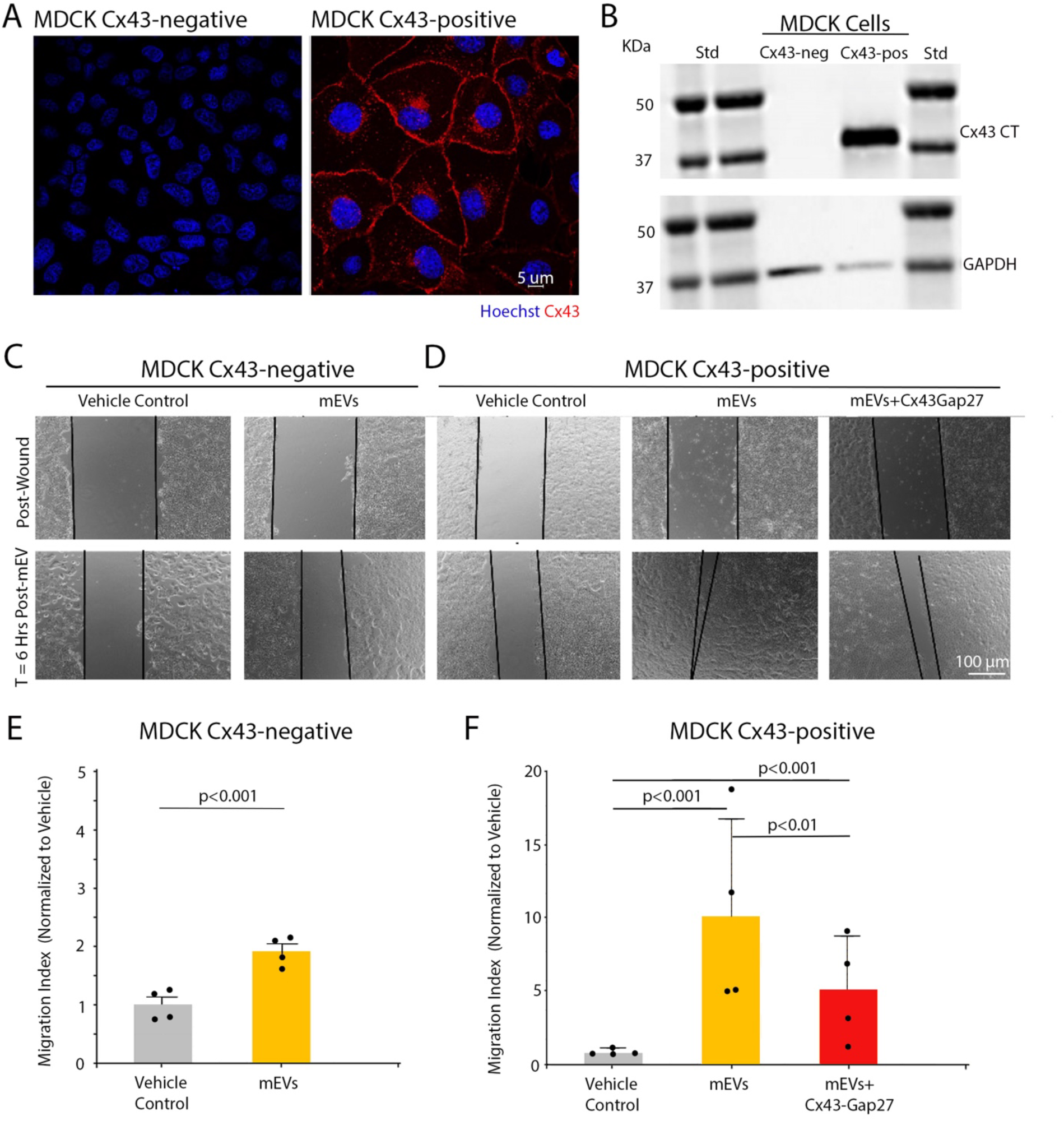
mEV-induced scratch wound healing in MDCK cells expressing (CX43+) and not expressing (Cx43-). A) Representative immunofluorescent images of MDCK Cx43-negative (Cx43-) and MDCK Cx43 positive cell (Cx43+) lines labeled with Cx43 CT antibody (against Cx43 CT amino acids 357-382) (B) Western blots of Cx43- and Cx43+ MDCK lysates confirm the presence and absence of the expected ∼Cx43 kDa band consistent with full-length protein, as detected by the Cx43 CT antibody. C) Cx43-MDCK monolayers immediately following scratch wounding (T=0, upper panels) and at T=6 hours of treatment with Vehicle Control or mEVs (lower panels), as indicated by labels. D) Cx43+ MDCK monolayers immediately following scratch wounding (upper panels) and at T=6 hours of treatment with Vehicle Control, mEVs or mEVs plus the Cx43 inhibitory peptide Gap27 (lower panels). Bar graphs for MDCK Cx43-scratch wound assays are shown in (E) and for Cx43+ cells in (F). Note that scratch wound re-growth in response to mEVs is much higher in the Cx43+ cells relative to controls and Cx43-MDCK cells. E and F) N = 4 experimental replicates for each group, error=SEM.

To investigate mechanism, we repeated mEV treatment of scratch wounded Cx43+ cells in the presence of Gap27 – a peptide mimicking the extracellular loop (EL) domain of Cx43 (Figs. 4D, F). Connexin EL domains mediate docking of connexin hemichannels (HC) and peptide mimetics of these domains block the pairing of HCs in closely apposed membranes – such as occurs during GJ channel formation ^37,38^ and between EVs interfacing with cell membranes ^16^. Consistent with a role for increased mEV-cell membrane interaction in Cx43+ cells, Gap27 (100 μM in culture media) significantly decreased (p<0.01), though did not abrogate, the effect of mEVs on Cx43+ cell migration following mechanical wounding (Fig. 4F).

### Cx43 Expression Affects Injury Dependent Cellular Uptake of mEVs In Vitro

The inhibitory effect of Gap27 on mEV-mediated upregulation of wound healing response of Cx43-expressing cells (e.g. Fig. 4) led us to explore effects of Cx43 expression on EV uptake. We fluorescently tagged mEVs with Cell Tracker Deep Red (CTDR), a dye that has been used in previous reports tracing EV uptake patterns in vitro and in vivo^39^. Similar to Calcein AM (e.g. Fig. 1D), esterified CTDR crosses the mEV membrane, but then becomes covalently linked to EV proteins, providing stable fluorescent labeling of the nanoparticle. We added CTDR-labeled mEVs to non-wounded Cx43- and Cx43+ MDCK monolayers, finding detectable, but relatively low numbers of fluorescent particles in cells from both lines (normalized to HOECHST-labeled nuclei), suggestive of moderate levels of mEV uptake at baseline (Fig. 5A upper and lower RH panels, 5B upper and lower RH panels). Scratch wounding (as shown in Figure 4) resulted in Cx43-negative MDCKs demonstrating a 2-fold increase (p<0.006) in CTDR particle counts (Fig. 5C). A significant increase (p<0.00001) in uptake of CTDR-labeled mEVs was also observed in scratch-wounded HDefs relative to unwounded monolayers of these cells (Supplemental Figure 2).

**Figure 5:**
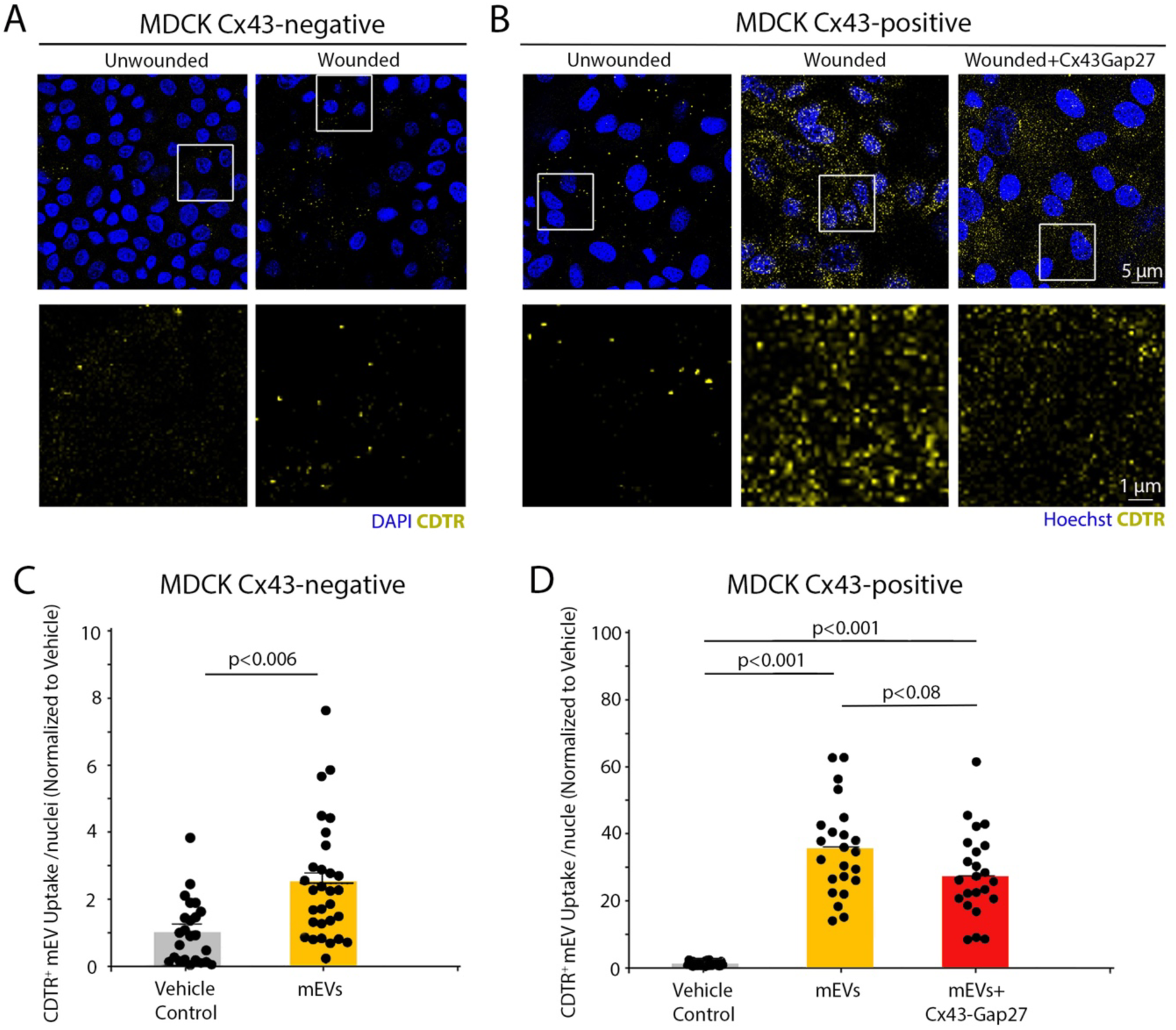
MDCK cells show increased uptake of mEVs following scratch wounding. A) Scratch wounding Cx43-negative (Cx43-) cells qualitatively indicates a modest increase in uptake of mEVs labeled with CTDR. B) Scratch wounding Cx43-positive cells (Cx43+) qualitatively indicates in a large increase in mEV uptake, which is partially abrogated by addition of exogenous 100 μM Gap27 to culture media. C) Particle count/nuclei quantification of mEV CTDR signals in MDCK Cx43-cells indicate a significant increase in mEV uptake in the wounded cultures. N=3 experimental replicates per group, each with 3 technical replicates, error=SEM. D) Particle count/nuclei quantification of mEVs taken up by MDCK Cx43+ cells shows a highly significant increase in mEV uptake in wounded cultures – at a rate over 10 times higher than that of unwounded Cx43+ cells. This effect is reduced by addition of exogenous 100 μM Gap27 to culture media, but not significantly so. N=3 experimental replicates per group, each with 3 or more technical replicates, error=SEM. CTDR counts from images are shown in (C) and (D) to illustrate range of variance.

**Supplemental Figure 2:**
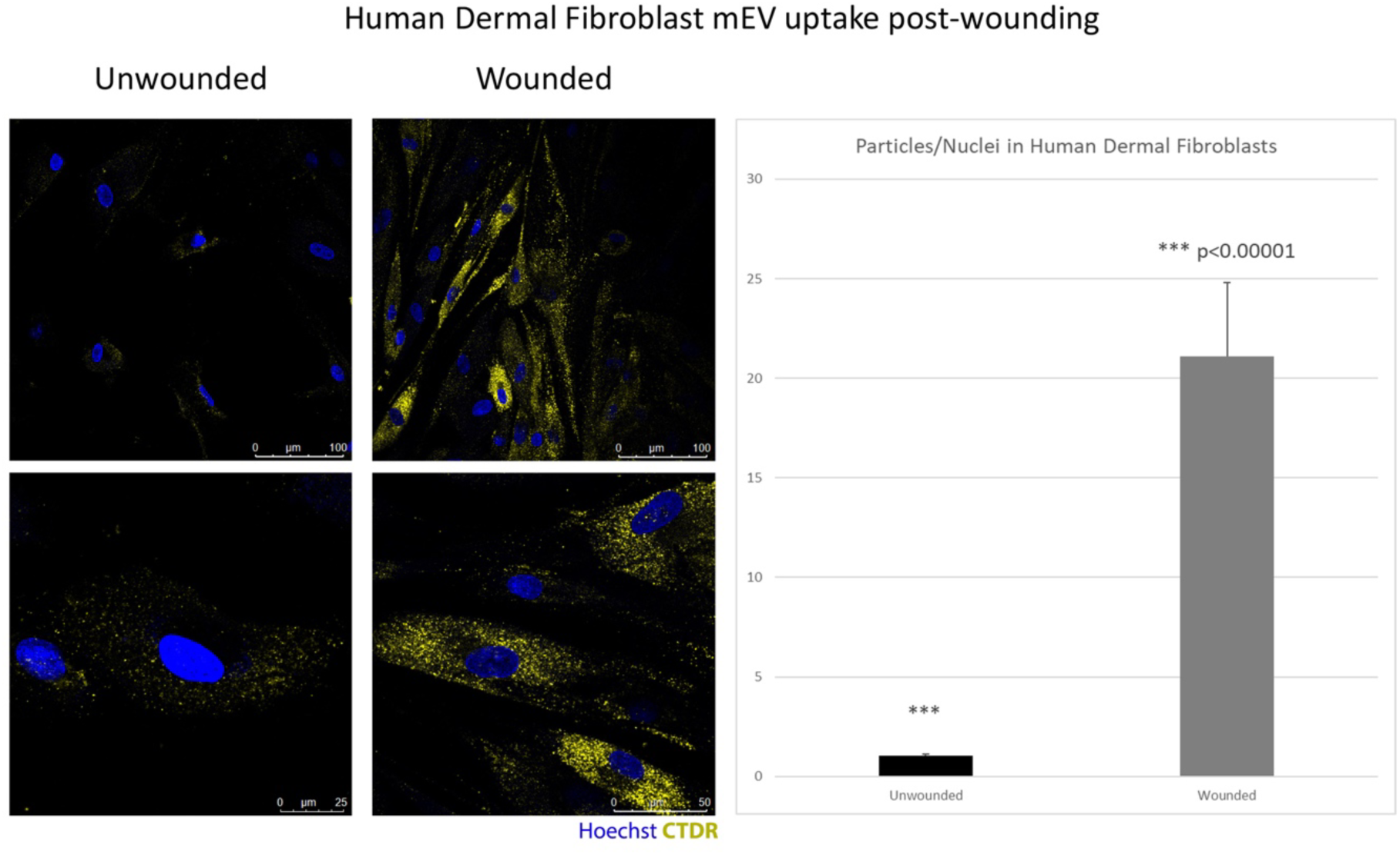
CTDR-labeled mEV uptake in unwounded and scratch-wounded monolayer cultures of human dermal fibroblasts (HDefs). LEFT) Unwounded HDefs exhibit modest levels of CTDR-labeled mEV uptake (yellow puncate signal). Fifteen minutes after scratch wounding, HDefs exhibit much higher levels of uptake of CTDR-labeled mEVs. RIGHT) Quantification of mEVs/nuclei in wounded vs. unwounded HDefs show a greater than 20-fold increase in punctate CTDR signals relative to unwounded cultures. N=3 experimental replicates per group. Error=SEM.

In contrast to Cx43-negative MDCKs, Cx43+ MDCKs showed a greater than 30-fold increase in punctate CTDR signals over unwounded Cx43+ MDCKs (p<0.001), indicating that Cx43 expression was likely associated with increased propensity to take up mEVs in response to scratch-wounding. Gap27 (100 μM) treatment resulted in a small, but not significant reduction (p=0.08) in mEV uptake induced by scratch wounding in Cx43+ MDCK monolayers (Fig. 5D). Interestingly, injury enhancement of uptake of mEVs was not just confined to cells proximal to the scratch wound in all the cell types studied: HDefs, MDCK Cx43-, and MDCK Cx43+ cells. Instead, fluorescent CTDR puncta occurred in cells throughout wounded monolayers.

### Cellular Uptake of Milk EVs is Acutely Increased in Injured Tissues in Vivo

We investigated the effects of tissue/cell wounding on mEV uptake using an in vivo model of cardiac ischemia reperfusion (I/R) injury ^40^. Left anterior descending coronary artery ligation was performed in CD1 mice to induce an I/R injury in the left ventricle (LV) (Fig. 6A). Immediately following surgery, CTDR-tagged mEVs (2 mg/kg body weight) were administered by oral gavage. Consistent with previous reports ^9^, mEV uptake into uninjured control hearts peaked over a 4-to 6-hour time course (Supplemental Fig. 3). In I/R injured LV four hours post-gavage, punctate CTDR signals were also observed – occurring at higher levels than in uninjured controls (Fig. 6B).

**Figure 6:**
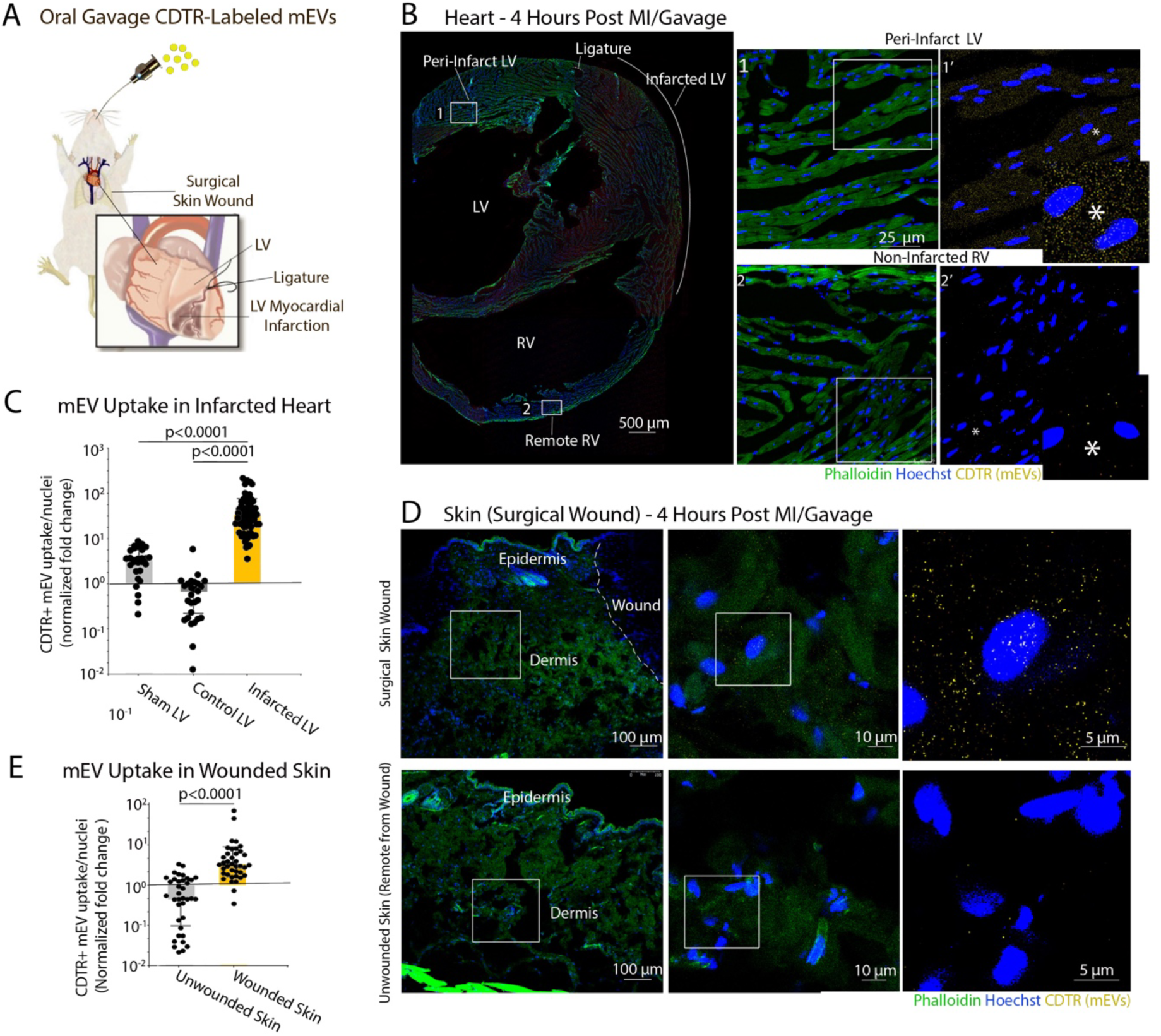
Skin and heart cells show increased uptake of mEVs following injury. A) Schematic of procedure by which CD1 mice underwent a surgical myocardial infarction resulting in an ischemia reperfusion (I/R) injury to the left ventricle (LV) and surgical wounding to skin. Directly following surgery mice were administered CTDR-tagged mEVs (2 mg mEVs/kg body weight) by oral gavage. B) Cryosection images of left (LV) and right ventricles (RV) 4 hours post-oral gavage of CTDR-labeled mEVs. The section is co-labeled with phalloidin-Alexa488 for actin (green) and Hoechst for nuclei (blue). The far left-hand image in (B) shows an overview of the ventricles, the point of ligature, infarcted LV, and RV myocardium are labeled. The upper right-hand panel (B1) shows a higher magnification of the boxed area adjacent the infarcted region of the LV in the overview shown in (B) with signals for CTDR (mEVs - yellow), actin (green) and Hoechst (blue nuclei). B1’ shows a higher magnification within the boxed region of B1 now only labeled for CTDR and HOECHST. High levels of CTDR puncta, consistent with mEV signals, can be observed in the cells in the LV. The lower right-hand panel (B2) shows a higher magnification of the boxed area in the non-infarcted RV also indicated in the (B) overview image. B2’ shows a higher magnification within the boxed region of B2. Only a few CTDR puncta can be observed in this region of non-infarcted RV. C) Particle count/nuclei quantification of mEVs taken up by heart cells in the infarcted LV show increases in mEV uptake relative to Sham surgical control LV (uninfarcted) and in the ventricles of uninjured control mice. N = 6 mice for each group, error=SEM. D) Cryosections of surgical skin wounds 4 hours post-oral gavage of CTDR-labeled mEVs of mice undergoing MI surgeries. The section is co-labeled by phalloidin-Alexa488 for actin (green) and Hoechst for nuclei (blue). D) As seen in heart tissues, dermal tissues adjacent wounds show increases in yellow CTDR puncta consistent with mEV-derived signals, relative to unwounded skin in the same mice remote from the injury. F) Puncta-particle per nuclei quantification of mEVs taken up by dermal cells indicates a significant increase in mEV uptake relative to remote uninjured skin. N = 6 mice for each group, error=SEM. CTDR counts from images are shown in (C) and (E) to illustrate range of variance.

**Supplemental Figure 3:**
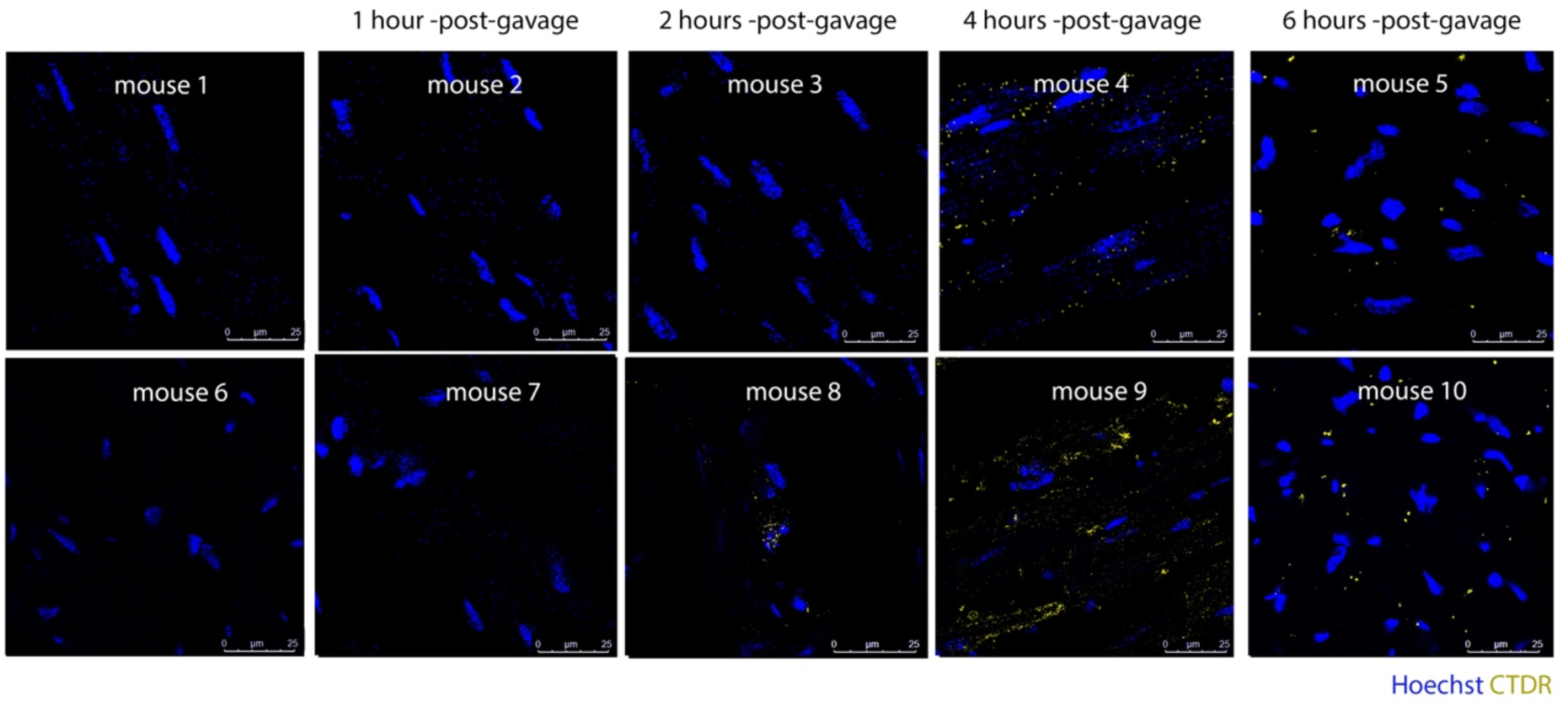
Fluorescent mEV-derived signals in ventricular myocardium of mice over a 6 hour time course following oral gavage of CTDR-labeled mEVs. Mice were provided with an oral gavage of 100 μL (2 mg/kg per mouse) CTDR-labeled mEVs, then were sacrificed at 1-, 2-, 4- and 6-hour time points following gavage. Hearts were excised and fixed, then in embedded in optimal cutting temperature (OCT) solution, sectioned and co-stained with Hoechst dye to delineate nuclei. mEV signals are shown in red, and can be seen to increase and peak at 4 hours, followed by reductions in signal at 6 hours.

Figure 6B shows this elevated punctate CTDR signal in the region of the LV adjacent to the infarct of an I/R injured heart, with signal increased in cardiomyocytes co-labeled with actin-phalloidin. CTDR signals are diminished in right ventricular myocardium, remote from the LV injury in this same heart (Fig. 6B). Whilst, variability in the counts of punctate CTDR signals were observed within both control and injured myocardial tissues, on average higher levels of fluorescent puncta (normalized to nuclei) occurred in the LV of infarcted hearts compared to uninjured controls (10-20-fold higher), as well as relative to Sham surgery controls (Fig. 6C). A visual survey of this elevated CTDR signal pattern and its variability across the left ventricle is shown in Supplemental Figure 4. This figure provides a montage of images taken from a cross-sectional view through the ventricle, providing a detailed visual representation of the phenomenon.

**Supplemental Figure 4:**
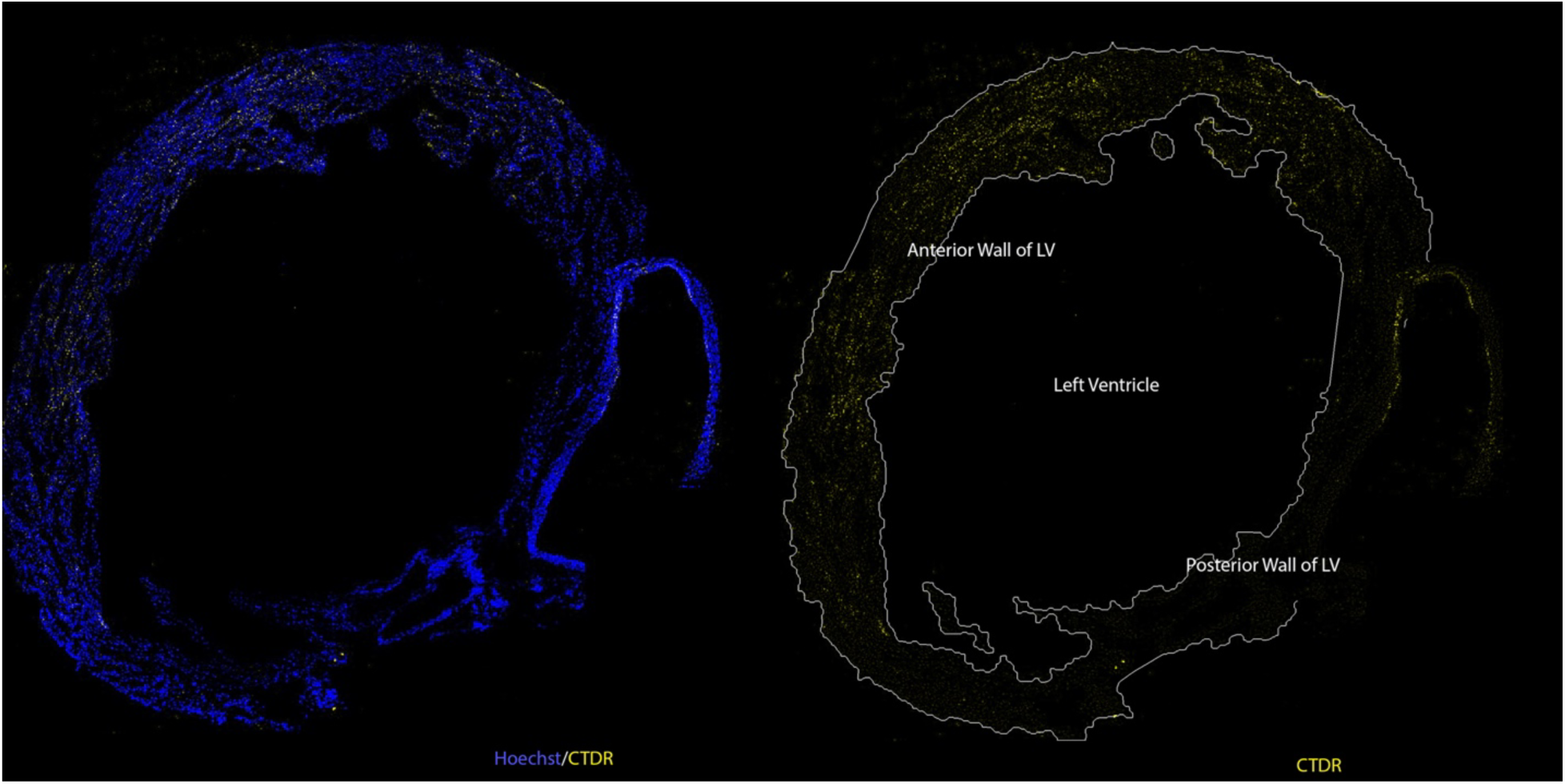
Survey view of LV of an infarcted mouse heart showing distribution of fluorescent mEV-derived signals. The mouse was provided an oral gavage of 100 μL (2 mg/kg per mouse) of CTDR-labeled mEVs just prior to myocardial infarction and the heart was sampled 4 hours after surgery. CTDR signal is broadly elevated in LV myocardium, though is lower distal in the posterior wall distal from the presumptive location of the I/R injury.

Interestingly, dermal tissues adjacent to surgical wounds on mice undergoing I/R injury also showed an average 2-3-fold increase in mEV uptake relative to unwounded controls within the same mouse (Fig. 6D, E). These findings, coupled with our in vitro scratch wound results (Fig. 5, Supplemental Fig. 2), suggest that injured cells exhibit enhanced uptake of mEVs relative to unwounded counterparts and that this property is maintained in vivo following oral ingestion.

### Development and Validation of the Approach to Loading Milk EVs with the Cx43 CT Mimetic αCT11

Given the increased targeting of mEVs to injured myocardial tissues (Fig. 6), as well as their bioactivity in vitro wound healing assays (Figs. 3 and 4), we sought to assess potential cardioprotective effects of native mEVs. We administered isolated concentrates of mEVs by oral gavage immediately prior to I/R injury and analyzed outcomes 24 hours later performing echocardiographic assessment of LV function and 2,3,5-triphenyltetrazolium chloride (TTC) staining of ventricular slices to assess infarct size (Supplemental Fig. 5). However, these initial studies indicated that mEVs showed little evidence of cardioprotective bioactivity, with no noticeable effects on infarct size or LV Ejection Fraction (%EF).

**Supplemental Figure 5:**
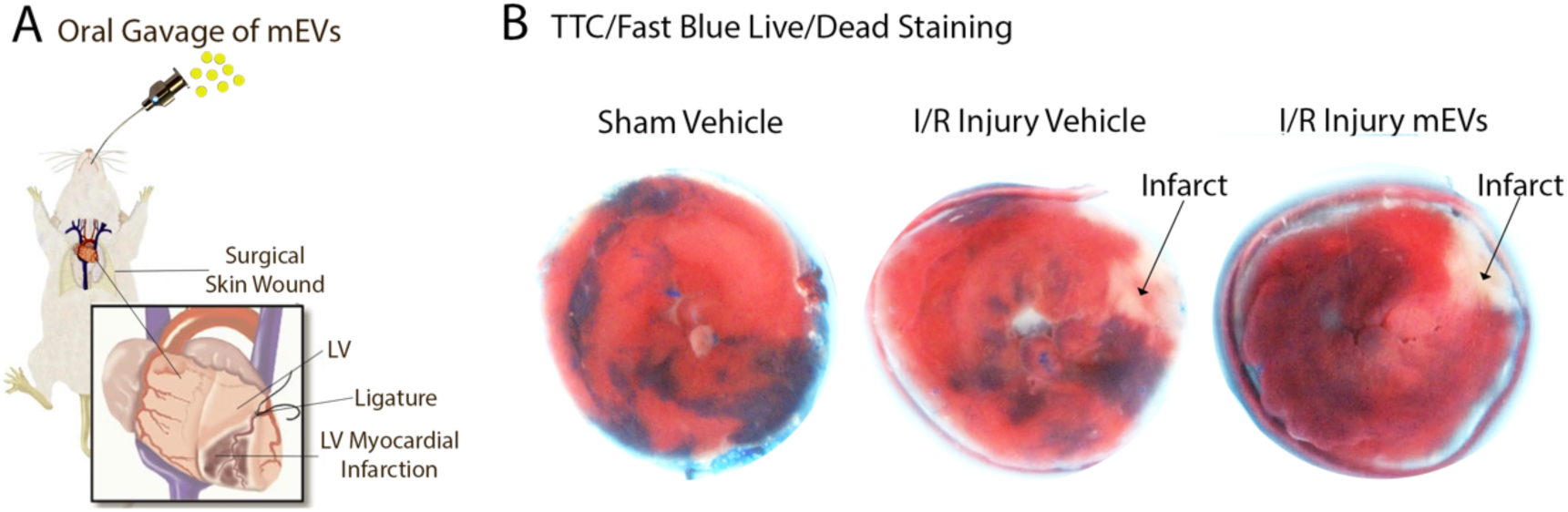
A) Schematic of procedure by which CD1 mice underwent a surgical myocardial infarction resulting in an ischemia reperfusion (I/R) injury to the left ventricle. Directly following surgery mice were administered a vehicle control solution or a solution containing an mEV concentrate (at 2 mg mEVs/per kg body weight per mouse) by oral gavage. B) Representative TTC/Fast blue stainings of 100 μm short-axis vibrotome slices through ventricles of Sham surgery, I/R injury cehicle control, and I/R injury mEV-treated mice 24 hours post-MI/mEV gavage. Note the infarcts in vehicle control and mEV-treated mice are of similar size.

It is has been reported that αCT11, a peptide mimetic of the Cx43 CT (RPRPDDLEI), is able to reduce ischemic injury size, preserve left ventricular function and reduce arrhythmogenic substrates in small and large animal ex vivo models of myocardial I/R injury ^22,27,28^. We thus sought to determine whether: 1) mEVs could be loaded with αCT11; and 2) if provision of small EVs containing this peptide provided acute cardioprotection at levels we have previously found for αCT11 (when not EV-encapsulated) in an ex vivo mouse model of cardiac I/R injury ^22^.

To enable EV loading, we used a strategy based on our finding that esterified Calcein AM is taken up and retained in mEVs (Fig 1D). We synthesized an esterified version of αCT11 in which the 4 carboxyl groups (Aspartic and Glutamic acid residues and the CT residue) were converted to methyl esters to produce a peptidic molecule referred to as 4OMe-αCT11 (Fig. 7A). Uptake of esterified αCT11 into mEVs was first assessed by Matrix-Assisted Laser-Desorption Ionization mass spectrometry (MALDI-MS, Fig. 7B-E). It was determined that whilst 4 methyl ester molecules dominated before loading into mEVs (Fig. 7D), following incubation of mEVs with 100 μM 4OMe-αCT11 the majority of peptide species detected by MALDI-MS within the nanovesicles comprised just 1 or 2 methyl esters (Fig. 7E). This suggested that as with Calcein AM, de-esterification within the EV was associated with retention of esterified αCT11 as a cargo molecule.

**Figure 7:**
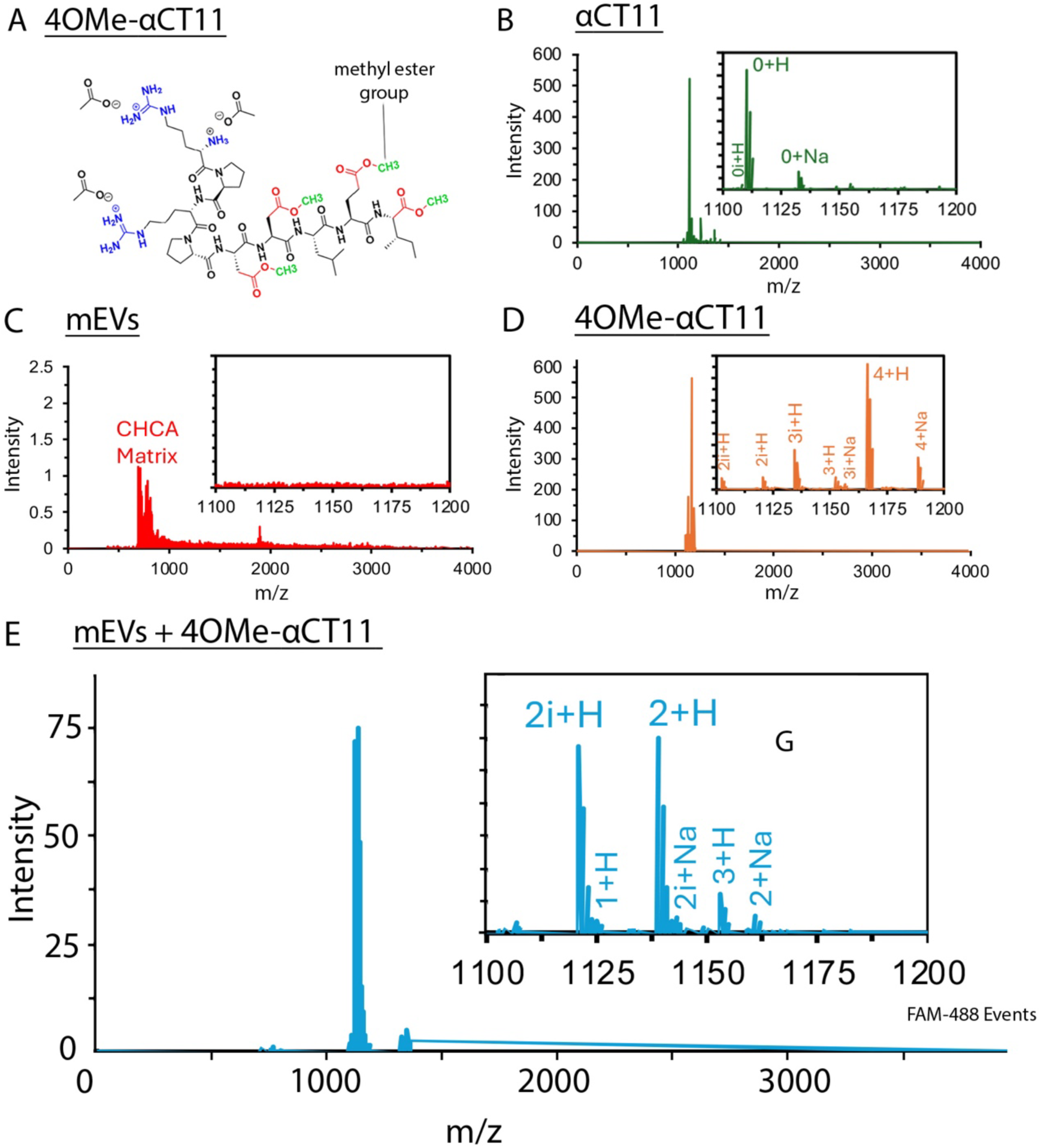
Mass spectrometry of esterified 4OMe-*α*CT11 prior to and after loading into mEVs: A) Schematic of 4OMe-αCT11 showing ester-linked methyl groups at location of where COOH groups occur in the natural, unesterified peptide. B-D)Matrix-assisted desorption ionization (MALDI) mass spectrometry (MS) of unesterified αCT11 (B), unloaded control mEVs (C), and 4OMe-αCT11 (D). E) MALDI-MS of mEVs loaded with 100 μM 4OMe-αCT11. Supplemental Table 1 provides a description of the esterified αCT11 species detected by MS in (B-E). Note that in panel (D), the dominant molecule is the 4 ester species, whereas in (E) after loading 4OMe-αCT11 into mEVs, the dominant molecules are the 2 ester species, consistent with the lability of aspartic acid esters we previously reported^33^.

**Supplemental Table 1:** Esterified αCT11 species detected by Matrix-assisted desorption ionization (MALDI) mass spectrometry in figure 7B through 7E, including information on numbers of ester and imide groups, co-ion and molecular mass for each molecule. Note that the 4 methyl ester+H (4+H) and 4 methyl ester++Na (4+Na) species were only detected in samples not co-incubated with mEVs.

| Label | αCT11 species | | Co-ion | Mass |
| --- | --- | --- | --- | --- |
|  | # of esters | # of imides |  | Da |
| 0+H | 0 | 0 | H | 1110.6 |
| 0+Na |  |  | Na | 1132.6 |
| 0i+H | 0 | 1 | H | 1092.6 |
| 1+H | 1 | 0 | H | 1124.6 |
| 2+H | 2 | 0 | H | 1138.6 |
| 2+Na |  |  | Na | 1160.6 |
| 2i+H | 2 | 1 | H | 1120.6 |
| 2i+Na |  |  | Na | 1142.6 |
| 2ii+H | 2 | 2 | H | 1102.6 |
| 2ii+Na |  |  | Na | 1124.6 |
| 3+H | 3 | 0 | H | 1152.6 |
| 3+Na |  |  | Na | 1174.6 |
| 3i+H | 3 | 1 | H | 1134.6 |
| 3i+Na |  |  | Na | 1156.6 |
| 4+H | 4 | 0 | H | 1166.6 |
| 4+Na |  |  | Na | 1188.6 |

To confirm loading of bioactive peptide into mEVs flow cytometry-based vesicle detection and the ECIS electric wound assay were used. First, we generated 6-carboxyfluorescein (FAM) tagged 4OMe-αCT11 and using flow cytometry, determining that fluor-conjugated peptide was taken up into mEV isolates in a dose-dependent manner (Fig. 8A-D). The highest concentration of fluorescent 4OMe-αCT11 tested, 100 μM, resulted in detectable signal in 85 % to 95 %, of vesicles (Fig. 8C,D). Second, using the electric wound assay it was determined that mEVs loaded with 100 μM 4OMe-αCT11 showed enhanced healing of electrical wounds in HDef monolayers compared to unloaded mEVs (p<0.00001) or αCT1 peptide. This elevated bioactivity, together with the evidence from mass spectrometry (Figs. 7B-E) and flow cytometry-based vesicle detection (Figs. 8A-D), confirmed efficient loading and bioactivity of mEV-encapsulated peptide.

**Figure 8:**
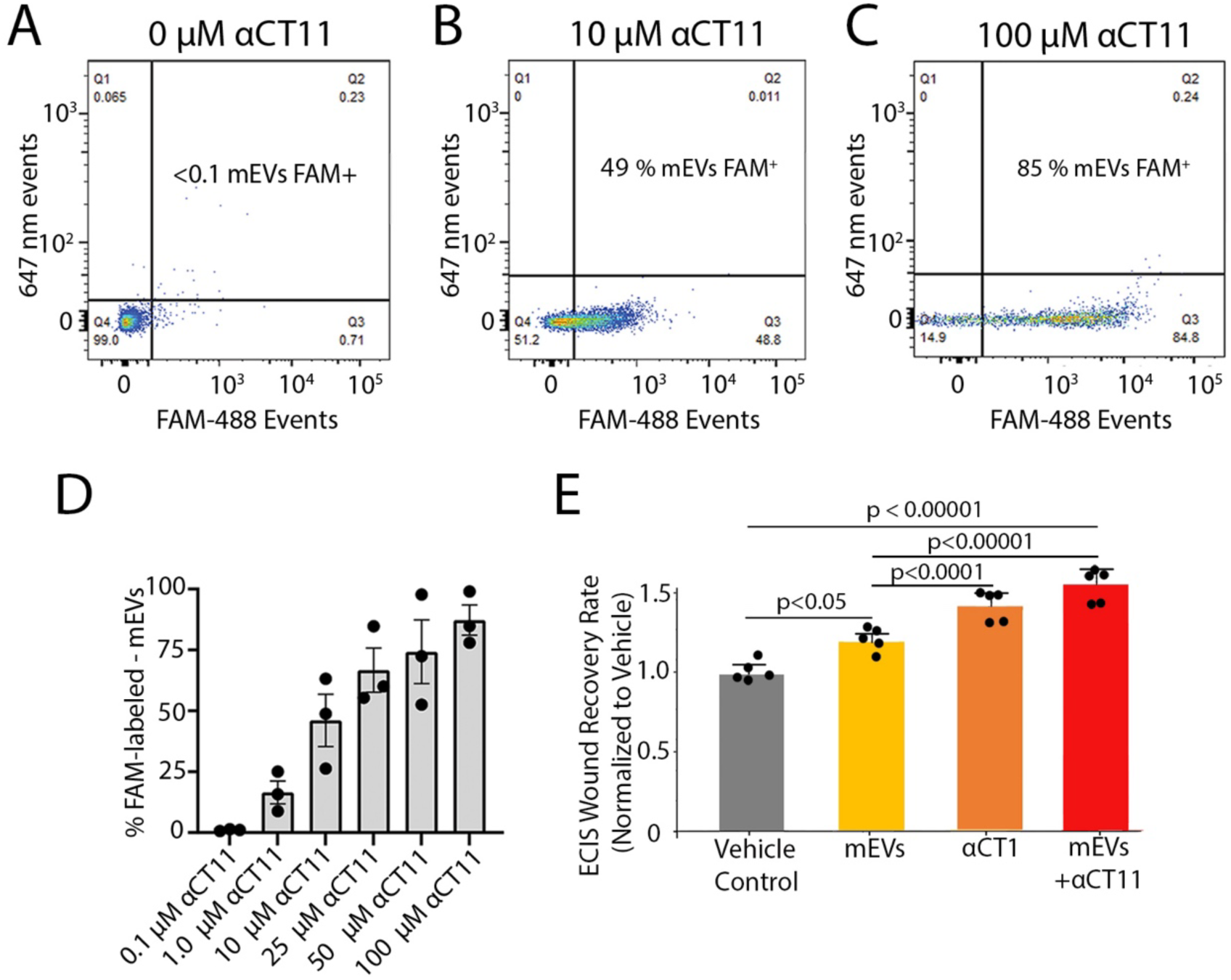
Assay of esterified-*α*CT11 loading into mEVs and bioactivity of peptide-loaded mEVs: A-D) Flow cytometry of mEVs loaded (or not peptide-loaded as in A) with different concentrations of fluorescent FAM-conjugated 4OMe-αCT11. Unloaded mEV controls were used to set the fluorescent gates and showed minimal background fluorescence (8A). mEVs incubated with 10 (8B) and 100 μM (8C) concentrations of FAM-conjugated 4OMe-αCT11 show increasing amounts of peptide uptake into mEVs. D) Flow cytometry quantification of uptake of fluorescent 4OMe-αCT11 over a concentration range of 0.1 to 100 μM. A mean of 85 % of nanoparticles are positive when mEV are incubated in the highest 100 μM concentration of fluorescent FAM-4OMe-αCT11. N = 3 per experimental group, error bars = SEM. E) Quantification of healing rates of HDefs in ECIS electric wound assays for vehicle control, unloaded mEVs, and 100 μM αCT1 (non-EV encapsulated) treatments, as well as mEVs loaded with 4OMe-αCT11 at 100 μM. Average healing rates after treating with unloaded mEVs, αCT1 and peptide-loaded mEVs are significantly elevated over that of the vehicle control group. Note that the 4OMe-αCT11-loaded mEVs exosomes accelerate healing to rates equal to or greater than 100 μM αCT1, which in turn are also both significantly increased over the unloaded mEV treatment. N = 4 per experimental group, error bars = SEM.

### αCT11-Loaded EVs Provide Cardioprotection when Administered before or after Myocardial Infarction

We next assessed the effects of peptide-loaded mEVs in an in vivo model of cardiac ischemia-reperfusion injury. mEVs loaded with 100 μM 4OMe-αCT11 were administered to mice via oral gavage at a dose of 2 mg/kg immediately prior to surgical induction of I/R injury to the left ventricle (LV). Morphological and functional outcomes were assessed by TTC staining of ventricular slices to quantify infarct size (Fig. 9A,B) and echocardiography (Fig. 9C, D) 24 hours after the myocardial infarction. Relative to vehicle and unloaded mEV controls, mEVs containing 4OMe-αCT11 resulted in significant improvements in LV ejection fraction following I/R injury (Fig. 9D). LV myocardial infarct size in mice treated with mEVs containing the exogenous Cx43 CT peptide (i.e., αCT11) demonstrated a greater than 60 % reduction (p<0.001) relative to injured vehicle controls (Fig. 9B). The infarct size in hearts receiving αCT11-loaded mEVs just prior to myocardial infarction was also significantly smaller than I/R injured mice receiving unloaded mEVs (p<0.004).

**Figure 9:**
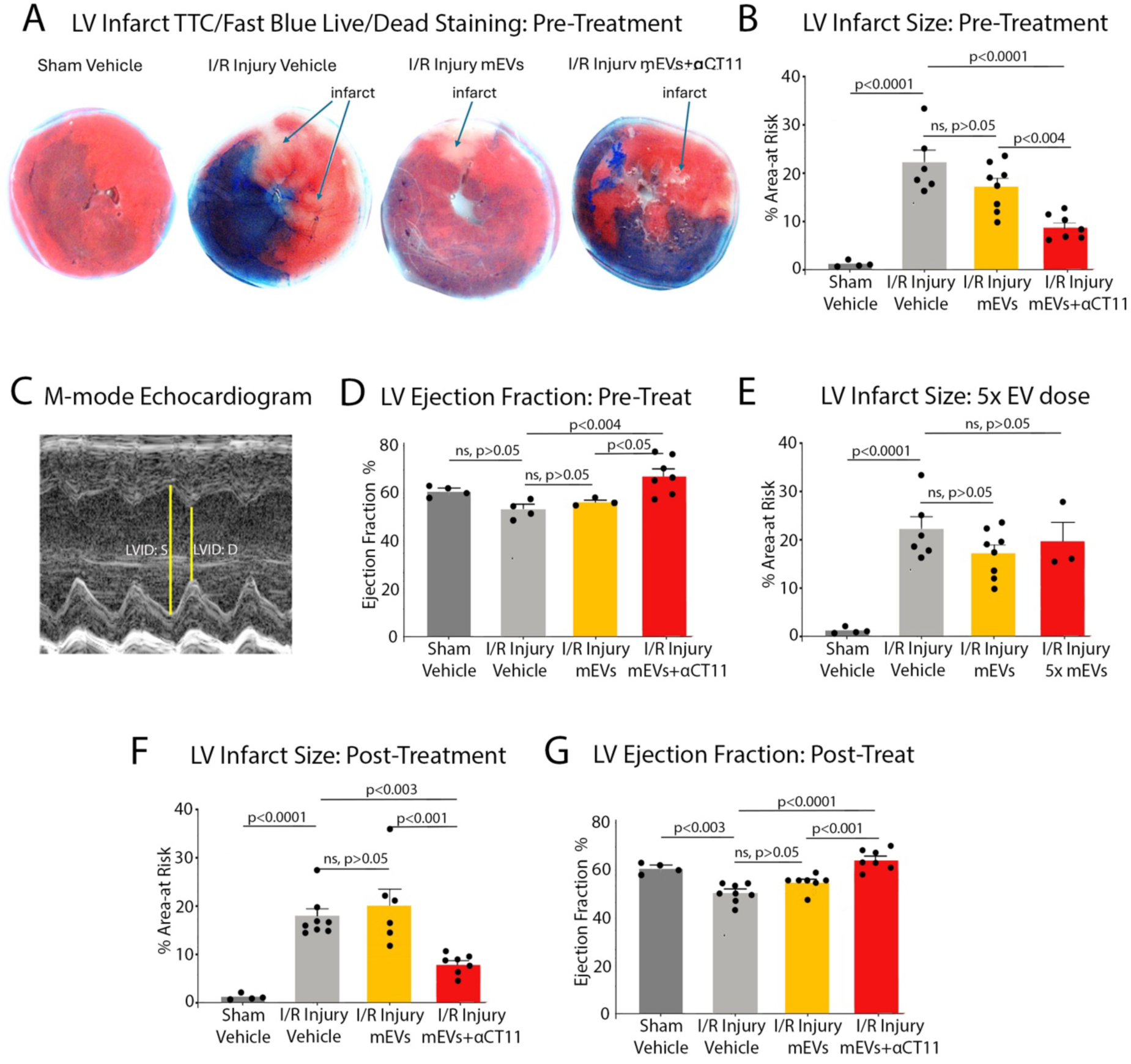
Pre- and post-infarction treatments with mEVs loaded with esterified *α*CT11 provide cardioprotection from ischemia-reperfusion injury to the heart in vivo. A) Representative TTC/Fast blue staining of 100 μm short-axis vibrotome slices through ventricles of Sham surgery control, I/R injury vehicle control, and I/R injury mEV- and 4OMe-αCT11-loaded mEV-treated mice 24 hours post-myocardial infarction. A single oral gavage of vehicle control or mEV (at 2mg/Kg per mouse) treatments of 100 (L were given just prior to the I/R injury anesthesia and surgery. B) TTC/Fast blue quantification of percentage (%) infarct area-at-risk in ventricular slices from the same 4 pre-infarction control and treatment groups as in (A) show significant increases in infarct size in I/R vehicle control and a modest, but not significant decrease in infarct size in mEV-treated mice, relative to I/R vehicle controls. Mice receiving 4OMe-(CT11-loaded mEVs (mEV+(CT11) demonstrate highly significant reductions in infarct size (a >60 % decrease) relative to I/R Injury vehicle controls. N=4-7 mice per group, error=SEM. C) Representative M-mode echocardiograph from a Sham surgery control mouse showing cyclic changes in LV internal diameter at systole (LVID:S) and diastole (LVID: D). LV ejection fraction (%EF) as measured from M-mode echocardiographs of the same pre-infarction control and treatment groups as in (A). Mice receiving 4OMe-αCT11-loaded mEVs demonstrated significant improvements in %EF relative to I/R injury vehicle and I/R injury mEV groups. N=3-6 mice per group, error=SEM. E) TTC/Fast blue quantification of percentage (%) infarct area-at-risk in ventricular slices from sham surgery control, I/R injury vehicle control, and I/R injury mEV groups as in (A). A fourth group of mice that received 500 μL of mEVs (i.e. a 5x dosage at 10 mg/Kg of mEVs per mouse) given just prior to I/R injury anesthesia and surgery. No decrease in infarct size was observed in mice receiving the 5x mEV dosage relative to I/R vehicle controls. F) TTC/Fast blue quantification of percentage (%) infarct area-at-risk in ventricular slices (N=4-7 mice per group, error=SEM) and G) % Ejection Fraction (N=4-7 mice/group, error=SEM) of analogous control and treatment groups as in (A), accept that oral gavage of controls and treatments with mEVs were given after surgical closure following induction of myocardial infarction. Mice receiving post-infarction 4OMe-αCT11 loaded-mEVs (mEV+αCT11) demonstrated a highly significant reduction in LV infarct size (∼60 % decrease) and LV ejection fraction relative to I/R Injury vehicle controls. infarct size and %EF in the group of mice receiving post-infarction treatment with native, unloaded mEVs did not differ significantly from I/R vehicle control mice.

Pre-infarction treatment of mice with native, unloaded mEVs did not provide a decrease in infarct size or improvement in LV function relative to vehicle controls (Figs. 9A-D). This being said, it was observed that this pre-treatment was associated with a small reduction in the size of the I/R wound and slight improvement in LV ejection fraction – albeit that these variables never reached statistical significance relative to vehicle controls (Fig. 9B, D). To further study whether native mEVs may show cardioprotective bioactivity, we provided mice with the equivalent of a 5-fold increase in dose, giving unloaded mEV concentrates by oral gavage at 10 mg/Kg over the usual 2 mg/Kg amount. However, this increased dose of native mEVs appeared not to have any detectable effect on I/R injury severity, with no significant reduction in myocardial infarct size below that of the 2 mg/Kg mEV dose or vehicle control hearts (Fig. 9E).

To be clinically useful to patients, such as those experiencing a myocardial infarction, a therapeutic would need to be given after an ischemic insult to the heart – i.e., after a myocardial infarction has been diagnosed. We thus carried out treatments of mice after I/R surgery – administering 4OMe-αCT11-loaded mEVs 10 minutes after surgical closure following completion of the 40-minute ischemic and 10-minute reperfusion phases of the injury model. Relative to vehicle and unloaded mEV controls, post-injury oral gavage with mEVs containing 4OMe-αCT11 (dosed at 2mg/Kg) provided decreases in LV infarct size similar in magnitude (∼60 %) to those measured in mice treated with peptide-mEVs prior to induction of myocardial infarction (Compare Figs. 9B,F). Similarly, peptide-loaded mEVs facilitated significant improvements in LV ejection fraction when provided by oral gavage after the I/R injury surgery relative to controls (Fig. 9G). As was the case in the pre-treatment paradigm, native, unloaded mEVs given post-myocardial infarction had no significant effect on either infarct size or LV function (Fig. 9F,G).

## Discussion

This study leverages the ability of mEVs to be preferentially taken up in injured cells to deliver therapeutic Cx43-derived peptides in a scalable manner suitable for oral administration. We further show that a subpopulation of mEVs contain Cx43, pointing to the potential biological cooperatively of Cx43 and mEVs in disease progression and cell-cell communication. Milk EVs increase cell migration in scratch-wounded monolayers of human dermal fibroblasts and MDCK cells in a Cx43-dependent manner, with uptake of mEVs into cells increasing with wounding in vitro and in vivo. Whilst native mEVs appear to be preferentially taken up by injured tissues, they do not significantly affect acute injury severity in a mouse ischemia reperfusion model of left ventricular injury. However, co-administration of mEVs loaded with an esterified mimetic of the Cx43 CT (4OMe-αCT11) results in a cardioprotective effect. This suggests that, although mEVs naturally contain potentially bioactive Cx43-related polypeptides, and exhibit increased uptake in injured tissues, their therapeutic potential in cardiac injury may be enhanced through strategic loading with bioactive Cx43-derived peptides.

The use of an esterified αCT11 peptide to load milk extracellular vesicles (mEVs) represents a technical advancement. This method provides a feasible way to load various small hydrophilic and negatively charged molecules into EVs, avoiding methods more adverse to nanovesicle integrity, like sonication or osmotic shock^41^. Our approach builds on key observations, including the uptake of esterified Calcein AM by mEVs, suggesting ester hydrolytic activity within them. The mechanism involves: 1) Converting negatively charged carboxyl groups in αCT11 to hydrophobic moieties; 2) Enhanced uptake due to these hydrophobic/charged-groups; and 3) Hydrolysis of ester bonds, which could lead to trapping of the peptide inside the mEV as it becomes less hydrophobic and/or more negatively charged during de-esterification. Optimal loading was achieved by converting available negatively charged side-chains on αCT11 to methyl esters. This comprehensive esterification maximizes the peptide’s ability to permeate the mEV membrane and ensures retention within the nanovesicles after ester loss.

We previously demonstrated that Cx43 CT peptides containing RPRPDDLEI (i.e., αCT11) reduce cardiac I/R injury severity in an ex vivo model^22^. This study shows that oral gavage of mEVs loaded with esterified αCT11 confers similar levels of cardioprotection in vivo when given acutely following I/R injury. Unprotected αCT11 peptide (i.e. non-EV encapsulated) rapidly degrades in blood serum and circulation, rendering it ineffective when injected intravenously in rodent I/R injury models ^26^. However, mEV-encapsulated αCT11 survives both circulatory breakdown and gastrointestinal uptake. This approach could enable targeted delivery of other therapeutic peptides using milk EVs as carriers, potentially improving oral bioavailability and efficacy of peptide-based treatments for various conditions beyond ischemic heart disease.

Emerging evidence suggests that connexins significantly influence the structure and function of extracellular vesicles (EVs) ^42^. Notably, EVs from connexin-expressing cells contain functional connexin hemichannels^16,43^. The detection of connexin 43 (Cx43) polypeptides in a subpopulation of EVs from cow milk supports these findings. However, the specific origin of this minority population of Cx43-positive nanovesicles remains uncertain. Potential sources include luminal breast epithelial cells, which express Cx43 ^44^, as well as breast myoepithelial cells^45^ and adipose tissues^46^, where Cx43 levels reportedly increase during parturition. Additionally, the possibility that EVs infiltrate milk from non-breast sources such as via the blood or lymph circulation cannot also be excluded.

A key finding is the identification of truncated CT Cx43 polypeptides in milk EVs. This builds on research showing that Cx43 CT isoforms can be generated through alternative translation of Cx43 mRNAs, with the main species being a 20 kDa molecule called GJA1-20K ^18,19^. Our experiments suggest a Cx43 CT peptide corresponding to this GJA1-20K isoform, along with other minor CT polypeptides of lower molecular mass. These may result from alternative translation ^47^ or proteolytic cleavage, as Cx43 is known to be susceptible to proteolysis ^48^. Future work could focus on amino acid sequencing of their N-termini to determine if naturally occurring Cx43 CT sequences in mEVs are more consistent with alternative translation or proteolytic cleavage.

The GJA1-20K-like Cx43 CT polypeptide we find in mEVs has been reported to affect cellular activity, including epithelial-mesenchymal transition ^49^, mitochondrial dynamics ^50^, actin organization, cell migration, and proliferation mitochondrial dynamics actin organization^51^. Exogenous GJA1-20K expression protects cardiac muscle against I/R injury ^20^ and reduces arrhythmic right ventricular cardiomyopathogenic disease ^21^. The last 9 amino acids of Cx43 (i.e., RPRPDDLEI) have been proposed as having significant roles in these bioactive properties of GJA1-20K ^52,53^. Synthetic peptides incorporating these 9 amino acids (e.g., αCT1 and αCT11) have been shown to accelerate wound closure in vitro and in vivo, as well as beneficially modulate injuries to eye and cardiac tissues in preclinical and clinical models^24,25^. To determine if the mEV effect on migration rates in wounded fibroblast and MDCK cell lines is due to these native Cx43 CT sequences, loss-of-function experiments on these polypeptides in mEVs, followed by evaluation of wounding response effects, are necessary.

Whilst our study demonstrated accelerated cell migration in wound assays in vitro, we did not observe significant cardioprotective effects of unloaded native mEVs in our in vivo I/R injury model. This discrepancy highlights the complexity of translating in vitro findings to in vivo applications and underscores the need for further studies of the bioactivity of mEVs in different physiological contexts. For instance, our ECIS results show that the mEV effect, though significant, is approximately half that of αCT1 peptide (i.e. non-mEV encapsulated αCT1). Thus, although milk-derived EVs suggest promise in regenerative healing scenarios, it may be that efficacy could be enhanced through targeted isolation of subpopulations or by focusing on specific bioactive components. For example, the modest effects seen in ECIS and I/R injury may be because only a minority of mEVs show evidence of the presence Cx43 and its bioactive CT isoforms. Future research should aim to elucidate the broad spectrum of mEV cargo molecules and their individual contributions to wound healing and tissue repair processes.

Consistent with effects observed in our in vitro wounding models, mEVs have shown potential in modulating injury and inflammatory processes across various tissues in vivo ^11,12,54^. Studies have highlighted their beneficial effects on injured gastrointestinal tract, heart, and skin ^55,56^. Reports suggest that bovine EVs from milk and colostrum may promote scar-free skin healing in mice ^57^, and reduce isoproterenol-induced cardiac fibrosis ^58^, respectively. Additionally, bovine milk-derived EVs have been reported to demonstrate promise in repairing UV-irradiated dermal cells ^59^. While most studies focus on microRNAs (miRNAs) as the likely primary mediators of mEV effects, our work, and that of others, suggest a more complex picture. Some EV populations may not incorporate substantive concentrations of miRNAs or effectively deliver these to recipient cells ^60,61^. Also, the less than 4-hour initial response time to mEVs observed in in vitro wound migration studies appears too swift to be consistent with a mechanism involving transcriptional inhibition.

One of the most notable biological effects observed was the increased cellular uptake of mEVs associated with wounding. This was particularly so in the heart, where a LV I/R injury led to a more than 10-fold increase in mEV-associated signals within 4 hours compared to uninjured controls. While increased uptake of other EV types in response to injury or inflammatory disease has been reported ^62^, this study is the first to characterize this effect for orally administered EVs. Two key questions arise as to this increase in cellular uptake of mEVs in response to injury: 1) What is its mechanism? and; 2) What is its biological purpose? Regarding the second question, the acute increase in uptake in injured and inflamed tissues might reflect mechanisms involved in maternal contributions to infant defense against infection. Some EVs appear to modulate inflammatory responses to infection ^63^, but more research is needed on mEV-specific behaviors in relation to tissue injury.

With respect to the mechanism of injury induced mEV uptake, a few inferences can be made. Factors effecting uptake of mEVs reported in the literature include variances due to route of administration, tissue barriers, organ and cell type^64^. For example, differences in mEV-internalization rates in mammalian cell lines can vary as much as an order magnitude ^65^. Our study reveals that injury-induced mEV uptake in cultured cells is rapid (within 15 minutes), is affected by Cx43 expression, and is broadly manifest within wounded cultures. These observations suggest a cell-intrinsic phenomenon, possibly involving diffusible factors. For example, we have found that treatment with apyrase, which hydrolyzes extracellular ATP, inhibits mEV uptake in in vitro ^66^ – suggesting ATP release by injured cells may be involved. Further, the increased uptake observed in injured tissues in vivo likely doesn’t occur via signaling gradients characteristic of immune cell injury homing responses ^67^. Instead, injured cells may either become modified to enhance uptake of circulating mEVs or release factors that locally modify mEVs, increasing their uptake likelihood.

The targeted uptake of small extracellular vesicles (EVs), particularly in pathological conditions, has garnered significant research interest^68^. While efforts have been made to modify EVs for selective uptake, such as tumor targeting, milk-derived EVs (mEVs) offer unique advantages. Their low immunogenicity and potential anti-inflammatory properties enhance their therapeutic utility^71,72^. Notably, our findings herein demonstrate that the intrinsic ability of injured cells to increase mEV uptake persists through oral ingestion and trafficking to injury sites. This in vivo targeting capability of unmodified mEVs, combined with our novel drug loading approach and the potential for large-scale production from milk, presents a promising foundation for a drug delivery platform that could have broad applicability. Furthermore, the observed cardioprotective efficacy of 4OMe-αCT11-loaded mEVs when administered after ischemia/reperfusion injury to the heart suggests significant potential for translation into clinical treatments for myocardial infarction. These findings open new avenues for EV-based therapeutics and warrant further investigation into their broader applications in regenerative medicine.

## Materials & Methods

### Isolation of small extracellular vesicles (EVs) from bovine milk

EV isolation protocols used were based on those previously reported by Marsh and co-workers^10^. In brief, raw bovine milk at 4°C was obtained from Homestead Creamery of Wirtz, VA after morning milking. Subsequent protocol steps, except the 1-hour 37°C EDTA treatment, were performed at 4°C. Milk was transferred to sterile polypropylene tubes (Thermo Scientific, 75007585) and centrifuged at 5000 rcf (Sorval Legend X1R centrifuge with Sorval TX-400 75003629 rotor) for 30 minutes. Fat was removed by whisking the creamy upper layer of supernatant (SN) with filter paper and decanting the remaining SN from the pellet. The centrifugation, whisking and SN decanting steps were then repeated. The resultant SN was transferred to 250 mL centrifuge tubes (Nalgene) and centrifuged at 14,500 rcf (Beckmann Coulter Avanti, J-26 XP centrifuge with JLA 16.25 rotor) for 60 minutes. SN was then decanted from the pellet and this SN was then centrifuged at 22,600 rcf for 60 minutes four consecutive times, with the pellet being discarded after each spin. This solution was then filtered through 0.45 μm and 0.22 μm filters (MilliporeSigma) in sequence and then treated with 30 mM EDTA at 37°C for 60 minutes. The EDTA-treated filtrate was then subject to cross-flow filtration using a Repligen Krosflo Tangential Flow Filtration (TFF) system with a 500 kDa MidiKros TFF Filter (Repligen). Once the TFF solution reached ∼20% of starting volume, the EV-containing solution was diafiltered in sterile, degassed 10x volume HEPES buffer (100 mM NaCl, 20 mM HEPES, 4 mM KCL, pH 7.5), aliquoted in 1 mL volumes and stored at -80°C. Prior to use, solutions containing isolated EV concentrates were thawed and separated on an IZON qEV original 70 nm sepharose column (IZON, 1006881), and collected manually in a 96-well plate, and peak EV fractions (fractions 7-9) were pooled for subsequent use and analyses, including Nanodrop and spectrophotometry.

### Gel Electrophoresis and Western blot (WB)

Gel Electrophoresis and WB analysis was performed as previously described^10^. Samples were separated by sodium dodecyl sulfate polyacrylamide gel electrophoresis (SDS-PAGE) and transferred to a PVDF (MilliporeSigma, IPFL00010) membrane. Membranes were blocked in EveryBlot Blocking Buffer (Bio-Rad Laboratories, cat. # 12010020) for 5 minutes at room temperature. Overnight primary antibody incubation was performed and primary antibodies were diluted in the blocking buffer as follows: CD81 (Cell Signaling Technology, 56039S, 1:1000) CD9 (Novus Biologicals, NB500-494, 1:1000), Calnexin (MilliporeSigma, AB2301, 1:5000), TSG101 (Bethyl Laboratories Inc. A303-506A, 1:5000), Casein (Abcam, Ab166596, 1:2000), Cx43 Cytoplasmic-Loop (Invitrogen, PA5-11632, 1:1000), and antibodies against the Cx43 CT (MilliporeSigma, C6219, 1:4000 and Abcam, Ab87645, 1:1000). Following washing, the membrane was incubated for 1 hour at room temperature with secondary antibodies diluted 1:20,000 for mouse (Jackson ImmunoResearch, 715-035-150) and 1:20,000 for rabbit (Southern Biotechnology, 4050-05) in blocking buffer. Proteins of interest were visualized by chemiluminescence using a Bio-Rad ChemiDoc MP imager.

### Cx43 Immunoprecipitation

Pellets of 500 μL of mEVs were lysed with 500 of RIPA buffer with protease inhibitor. EV lysates were precleared by incubating with 50 μL of anti-rabbit IgG beads (MilliporeSigma, A8914) for 10 min on ice. Precleared lysates were then incubated overnight with 5 μg Cx43 CT antibody against an epitope corresponding to AAs 363 to 382 of the protein (MilliporeSigma, C6219). 50 μl of anti-rabbit IgG beads were added the next morning and solutions were then incubated at 4° for 2 hours. After incubation sample was centrifuged at 10000xg for 1 minute and the supernatant removed. The pellet was then washed three times with RIPA buffer, resuspended in Laemmli buffer and denatured at 95°C for 5 minutes. The sample was then centrifuged at 10000xg for 1 minute and the supernatant collected for immunoblotting using a second Cx43 CT antibody against an epitope corresponding to AAs 337 to 346 of the protein (Abcam, Ab87645, 1:1000), followed by detection as in the western blotting protocol.

### Flow cytometry-based vesicle detection

For detection of Cx43 in mEVs, the intracellular fixation and permeabilization buffer set (eBioscience) was used to fix and permeabilize mEVs for intravesicular staining. Pellets of 250 μL of mEVs were fixed by adding 100 μL of fixation buffer and incubated in the dark at RT for 20 minutes. After incubation, the samples were washed twice with 1 ml of permeabilization buffer and resuspended in 100 μL of the same buffer. The samples were then incubated overnight with either Cx43 Cytoplasmic-Loop (Invitrogen, PA5-11632), or Cx43 CT (MilliporeSigma, C6219) antibodies directly conjugated to Alexa-488 dye, using the FlexAble antibody labeling kit (Proteintech). After incubation, the labeled mEV samples were washed twice with 1 ml of permeabilization buffer and resuspended in 200 μL of PEB buffer (PBS, 5mM EDTA, and 0.5% BSA) followed by flow cytometry-based vesicle detection. Samples were collected on a BD Biosciences FACS Fusion Flow Cytometer. EVs were displayed on an FSC-A and SSC-A dot plot using log scale. A gate was drawn around the EVs and a subsequent plot of FSC-H (log) vs FSC-W (lin) was displayed and gated to remove aggregate events. Acquisition was set to record 10,000 singlet events per sample. Alexa 488 was detected with a 488nm laser excitation and a 530/30 bandpass filter. After acquisition the same gating methods were employed to analyze the samples using FlowJo software VX. To measure the loading efficiency of 4OMe-αCT11 using flow cytometry, we generated 6-Carboxyfluorescein (FAM) tagged 4OMe-αCT11. 250 μL of mEVs were incubated with FAM-tagged 4OMe-αCT11 for 3 hours at room temperature at concentrations indicated in results. After incubation, the samples were washed twice with 1 ml of PBS buffer and resuspended in 100 μL of PBS followed by flow cytometry-based vesicle detection, as described above for Cx43 in mEVs using flow cytometry.

### Nanoparticle Tracking Analysis (NTA)

NTA was performed on a NanoSight NS300 (Malvern Panalytical) at 20°C. EV concentrates obtained post-SEC were diluted 1:10 in HEPES buffer, sonicated in a Branson 2510 bath sonicator for 30 seconds. EV samples were then diluted (1:1,000 to 1:10,000 depending on sample) and loaded into the NanoSight low volume flow cell. Each sample was analyzed using a 405 nm laser with 5 consecutive 1-minute recordings at a constant flow of 10, as per Malvern software. Videos were analyzed using NTA software (Version 3.4) and nanoparticle counts and dimensions generated as we have previously reported ^10^.

### Transmission Electron Microscopy (TEM)

Prior to sample loading formvar coated copper grids (Electron Microscopy Services, FCF200-CU-50) were glow discharged using a Pelco Easy Glo (Ted Pella). For each following step, solution volume was 10 μL, and solution was removed by gently wicking with a small square of Whatman filter paper #1. Grids were coated with 0.01% lysine for 60 seconds, then rinsed twice in 10 μμL ddi water from a Millipore unit. Samples were loaded onto lysine coated grids for 10 minutes, then counterstained with uranlyess (Electron Microscopy Services, 22409-20). Samples were stored in a standard grid box at room temperature prior to TEM imaging.

### Laser Scanning Confocal Microscopy (LSCM) of Fluorescent Dye Loading

For both in vitro and in vivo tracking analysis, mEV samples were loaded with Calcein AM (Invitrogen, C1430) or cell tracker deep red (CTDR) (Invitrogen, C34565) at 20 μM for 2 hours at 37°C, then centrifuged at 16,873xg for 1 hour at 4°C to remove unincorporated dye. Supernatants were discarded, and pellets were resuspended overnight at 4°C prior to trituration. Samples were sterile-filtered prior to being administered in both scratch wound healing assays in vitro and by oral gavage in animal experiments, as described in a following section. Cells or tissues were also processed as described below, then imaged on a Leica TCS SP8 LSCM.

### Single Molecule Localization Microscopy

Single Molecule Localization Microscopy (SMLM) was performed in collaboration with the New York Institute of Technology (NYIT) and Nanometrix ^74^. The EV Profiler Kit (Oxford Nanoimaging) containing antibodies against the tetraspanins - CD9, CD63, and CD81 was used according to the manufacturer’s instructions. Briefly, a 2.5 µl of sample was gently mixed with 3.5 µL blocking buffer and allowed to incubate for 5 minutes prior to the addition of fluorophore-conjugated antibodies. In addition to antibodies provided in the kit, the following stains were used: goat anti-Connexin 43 (Abcam, ab219493) conjugated to Alexa Fluor 647 using the APEX Antibody Labeling Kit (ThermoFisher); and Alexa Fluor 488-conjugated wheat germ agglutinin (ThermoFisher). When necessary, PBS wash buffer was used to reach the final sample volume of 9 µl. Samples were allowed to incubate overnight at 4°C protected from light. The following day, samples, buffers and the chip were brought to room temperature. Immediately after surface preparation of the chip, the samples were mixed with the EV capture solution and loaded into the lanes on the chip. The chip with samples was incubated for 15 minutes protected from light, whereafter wash and fixation steps were performed. Immediately prior to imaging, freshly mixed Bcubed buffer was added to the lanes. Imaging was then performed and Nanometrix software was used for analysis and data output.

#### 4OMe-αCT11 Synthesis

4OMe-αCT11 was prepared as described previously,^32^ using Fmoc-solid phase peptide synthesis with a CEM Liberty Blue automated microwave-assisted peptide synthesizer. We selected 2-chloritrityl chloride as the resin (0.6 mmol/g) to furnish peptides with a COOH C-terminus (rather than an amide group). The first amino acid was coupled to the resin using potassium iodide (0.125 M in N, N-dimethylformamide (DMF)) and N,N-diisopropylethylamine (1 M in DMF), and the remaining amino acids were coupled using diisopropyl carbodiimide (1 M in DMF) and Oxyma Pure (1 M in DMF). Resin was treated with piperidine (20% v/v in DMF) to deprotect Fmoc groups preceding amino acid additions. After synthesis, peptide chains were cleaved from the resin using a deprotection cocktail composed of trifluoroacetic acid, reverse osmosis-purified H_2_O, triisopropyl silane, and 2,2-(ethylenedioxy)diethanethiol (92.5/2.5/2.5/2.5 v/v) for 3 h at room temperature under constant stirring. Deprotected peptides were separated from resin by filtration, then isolated by precipitation into diethyl ether and centrifugation (5 min, 2420 xg, 4 °C). The supernatant was discarded, and the peptide was washed again with diethyl ether and isolated by centrifugation under the same conditions. Peptide pellets were dried under vacuum for 1 h, dissolved in 5% (v/v) acetonitrile in H_2_O (both with 0.1% TFA) and frozen in liquid N_2_ immediately prior to lyophilization for 48 h. To esterify αCT11, methanol (1600 : 1 mol/mol) containing 5% HCl (v/v) was added to solid lyophilized peptide. The solution was stirred for 24 h at room temperature, and the esterified peptides were isolated and dried as described above. All crude peptides were purified using preparative RP-HPLC on a Waters Empower system equipped with a XBridge® Prep C18 optimum bed density chromatographic separation column (30 mm × 150 mm, 5 μm beads) and a photodiode array detector (Waters 2489 UV/visible) to achieve > 85% purity. Finally, to replace TFA counterions with acetate counterions, 4OMe-αCT11 was dissolved into 5% (v/v) acetonitrile in H_2_O (both with 0.1% TFA) and dialyzed against acetic acid, with 4 external solution changes. Peptide samples were first dialyzed against 30% (v/v) acetic acid in RO H_2_O for 4 h to replace the counterions, followed by three switches of the external solution to 0.1% (v/v) acetic acid, which were necessary to prepare the sample for lyophilization. After lyophilization, counterion-exchanged samples were aliquoted into 1 mg samples and lyophilized for storage as solids until use.

### Matrix-Assisted Laser Desorption/Ionization-Time of Flight (MALDI-TOF) Spectrometry

MALDI-TOF was measured using a Shimadzu MALDI-8030 mass spectrometer with a 200 Hz solid-state laser (355 nm). The instrument was calibrated with a standard MALDI calibration kit (TOFMix and CHCA matrix, Shimadzu) within 0.5 Da, for which samples were dissolved at 670 femtomoles per μL in 70% v/v ACN with 0.1% v/v TFA. Prior to measurement, 300 µg of either unloaded mEVs or 100 μM 4OMe-αCT11-loaded mEVs were dissolved in 0.25 mL of 94.95/4.95/0.10 H2O/acetonitrile (ACN)/trifluoroacetic acid (TFA) (v/v), sonicated for 5 min, and then filtered through 0.45 µm filters. For peptide standards, 170 µg of either αCT11 or 4OMe-αCT11 were dissolved and then underwent the same treatment. EVs and peptide standards were then co-crystallized in a 1:1 ratio (v/v) with a α-cyano-4-hydroxycinnamic acid (CHCA) matrix solution (5 mg mL−1 in 70% v/v ACN with 0.1% v/v TFA). Peaks with less than 2% intensity or those that could be attributed to a CHCA blank were not reported.

### Cell Culture

Cells used were human dermal fibroblasts (HDef; ATCC, PCS-201-012), as well as two lineages of Madin Darby Canine Kidney (MDCK) Cells-one line heterologously expressing high levels of Cx43 (Cx43+), and the other line with no detectable Cx43 expression (Cx43-) generously provided by Dr Paul Lampe (Fred Hutchison Cancer Center, Seattle, WA). Medium used for HDef was DMEM HG (4.5 g/L Glucose) with 2% Normal Calf Serum (NCS; Thermo Fisher/Gibco, 16010-159, Lot 2490415) and 4% Fetal Bovine Serum (FBS; Thermo Fisher/Gibco, 26140-079). Medium used for the Cx43+ MDCK cells was M199 (Millipore Sigma M4530) supplemented with 10% FBS and 1% Hygromycin B (Sigma H0654), while Cx43-MDCKs used M199 with 10% FBS and 1:125 Sodium Pyruvate (Invitrogen 11360-070). Cells were expanded and stored in liquid nitrogen until plating on standard polypropylene culture dishes in culture medium, as described above. Cells were expanded to confluency, then passaged into 12-well plates for analysis.

### In Vitro Scratch Wound Assay

For scratch wound studies in vitro, we used our previously published approach ^34^.^34^. Briefly, cells were plated in 12-well plates and given 2 days to adhere and grow prior to replacing with media not containg serum. Following serum removal, treatments as reported in results were carried out, i.e., mEVs at 100 μL of 100 μg/ml of mEVs loaded with 100 μM αCT11, vehicle control, and Gap27 (Apex Bio, HY-P0139) at 100 μM in media. Assay was performed 24 hours after treatment, using a 200 μL sterile pipette tip to scratch the surface of the well. Cells were then rinsed 1x in PBS and provided fresh culture medium supplemented with fresh control and treatment solutions. Cells were then imaged along the length of each scratch and images analyzed to determine initial scratch area. After 6 hours, cultures were rinsed in 1x PBS, fixed in 2% paraformaldehyde, then rinsed 4 times in PBS and stained in 1:20,000 Hoechst prior to imaging on a Leica SP8. Final and initial scratch areas were determined using automated ImageJ Area Analysis software abd migration was calculated by (Initial Area-Final Area)/Initial Area), with a larger number indicating greater movement towards a final area of 0. Estimates were further normalized to the average of the vehicle control group. Investigators performing the quantitation were blinded to treatment.

### ECIS In Vitro Wound Assay

Human Dermal Fibroblasts were expanded as described in “Cell culture and In Vitro Scratch Wound Assay” sections above. Monolayer confluency was measured in real time with an electric cell-substrate impedance sensing (ECIS) instrument (Applied Biophysics) to monitor migratory response. Briefly, 8W1E electrode ECIS arrays (250 µm diameter) were coated with 10mM L-cysteine, followed by 1% gelatin according to manufacturer guidelines. Electrodes were then stabilized in HDef complete media, and subsequently seeded with HDef cells at a density of 1×10^6^ cells/well in a final volume of 400 µL. Cells were incubated for 8-10 hours to achieve at least 80% confluency, which was confirmed by previously performed growth-phase profiles. 8 hours before electric wound induction, media was changed to HDef complete media supplemented with 10 µg/mL mitomycin C to inhibit cell proliferation. A 25 kHz, 2600 µA, 20 second electric wound was then applied to the cell monolayer and impedance was monitored until migration over the wounded area was complete. Cell migration completion time (t_heal_) was determined by a custom Python script that tracked changes in impedance over time. The impedance of cell-covered electrodes was measured with the multiple frequencies over time (MFT) mode at eleven frequencies between 62.5 to 64000 Hz (62.5, 125, 250, 500, 1000, 2000, 4000, 8000,16000, 32000, 64000). Investigators performing the quantitation were blinded to the treatment.

### Assay of mEV Uptake in Vitro

For uptake studies in vitro, cells were plated onto sterile coverslips within the plate, then given 2 days to adhere and grow prior to scratch assay. Assay was performed by using a 200 µL sterile pipette tip to gently scratch (or not) the surface of the cells, then cells were rinsed 1x in PBS (Invitrogen, 14080055) and provided fresh culture medium supplemented with 100 μL of 100 μg/ml of CTDR-labeled mEVs prepared as for the “LCSM” section. Some cells were also supplemented with Gap27 (Apex Bio, HY-P0139) at 100 μM in the media. Cells were given 15 minutes to take up mEVs, then were rinsed in 1x PBS and fixed in 2% Paraformaldehyde (Fisher O4042-500). Cells were then rinsed 4x in PBS, and stained with Hoechst dye (1:20,000, Life Technologies, H3569) and rinsed one additional time in PBS. Coverslips were then removed and adhered to microscope slides, and CTDR (red) and nuclear Hoechst (blue) signals were imaged on a Leica SP8 LSCM. Images were analyzed in ImageJ by saving the red and blue channels, converting to 8-bit, thresholding, and counting CTDR particle numbers and nuclei. Counts were done in triplicate with 10 images taken of each sample for an individual experiment n=30, with experiments being done in triplicate. Investigators performing the quantitation were blinded to the treatment.

### Myocardial Infarction and Skin Wound Healing Studies In Vivo

Adult male outbred ICR (CD-1) mice were supplied by ENVIGO (Indianapolis, IN). All animal experiments were conducted under the guidelines on humane use and care of laboratory animals for biomedical research published by the National Institutes of Health (No. 686-23, revised 20112011). The study protocol was approved by the Virginia Commonwealth University Institutional Animal Care and Use Committee. Mice received 100 μL or 500 μL mEV solutions at 2mg/Kg or the same volume of control solutions by oral gavage using a round-tip gavage needle, just before being intubated or immediately after myocardial infarction (MI) surgery, as described in the results. Experimental MI was induced as we have previously described ^40^, by a transient occlusion of the left descending coronary artery for 40 minutes followed by reperfusion for 10 minutes. Animals were anesthetized using isoflurane (1.5-3.0% in 100% O_2_) intubated orotracheally and ventilated on a positive-pressure ventilator. The tidal volume was set at 0.2 mL, and the respiratory rate was adjusted to 133 cycles/min. A left thoracotomy was performed at the fourth intercostal space, and the pericardium was stripped to expose the heart. The left descending coronary artery was then identified through a surgical microscope (Leica F40; Leica Microsystems, Wetzlar, Germany) and ligated with a 7.0 silk ligature, which was then released after 40 minutes to allow for reperfusion. A sham surgery (control) was performed by performing all the steps, except the transient occlusion of the artery. The mEVs, unloaded or loaded or with the 4OMe-αCT11 were administered through oral gavage at just prior to intubation and surgery or just after surgical MI as indicated in the results. At 24 hours following MI, animals underwent transthoracic echocardiography using a Scintica Prospect T1 imaging system (Scintica Instrumentation Inc, London, Ontario, Canada) and a 30-50 MHz probe. The heart was first imaged in the two-dimensional mode in the short-axis view of the left ventricle according to the American Society of Echocardiography recommendations, at the level of the papillary muscles below the mitral valve tip. The M-mode cursor was positioned perpendicular to the anterior and posterior wall to measure LV end-diastolic diameter (LVEDD), LV end-systolic diameters (LVESD), to calculate the ejection fraction using the Teichholz formula. Mice were then euthanized and infarct size measurement was performed using triphenyl tetrazolium chloride as previously described ^40,75^. The areas of infarcted tissue, the risk zone, and the whole LV were determined by computer morphometry using Image Pro Plus 6.0 software (Media Cybernetics, Silver Spring, MD). The investigators performing and reading the echocardiogram and TTC results were blinded to the treatment.

### Assay of mEV uptake in Response to Injury In Vivo

CD1 mice were administered 100 μL of CTDR-tagged mEVs by oral gavage at 2mg/Kg, then subject to MI surgery as in the preceding section. Mice were sacrificed at 4 hours post-injury, hearts and skin specimens were then immediately collected, including from the surgical skin wound tissues, as well as unwounded skin from the opposite side of the animal (“unwounded skin”). Tissues were then fixed in 4% paraformaldehyde and embedded in OCT (Fisher, 23-730-571) for cryosectioning on a Leica CM1850 UV onto Superfrost Plus Micro Slides (VWR, 48311-703). Sections were then coverslipped and imaged on a Leica SP8 LSCM. Images were analyzed in ImageJ by saving the individual red channel and converting to 8-bit, thresholding, and counting particle numbers. Samples were analyzed in triplicates with individual experiments also being performed in triplicate. Control groups (sham surgery) were supplied with equivalent volumes of free CTDR dye, and unloaded mEV’s were performed and showed only background signal (data not shown). Investigators performing the quantitation were blinded to treatment.

### Statistical analysis

Data is presented as mean ± SEM unless specified. Statistical analysis was performed using one-way ANOVA with post-hoc correction for multiple comparisons. Owing to heteroscedasticity of data in Fig. 4F, data were log transformed, whereafter it was found to conform to uniform variance by Bartlett’s test. P values on Fig. 4B are post-hoc corrected values following one-way ANOVA on the log transformed data. All statistical analysis was performed using GraphPad Prism v.10.0.2. A p<0.05 was considered statistically significant.

## Acknowledgments

The authors thanks Melissa Makris and Jaya Asseervatham for technical assistance. The authors acknowledge Dr. Lars Udo-Bellner and Amanda Charest for assistance with SMLM as part of the NYIT Imaging Center.

## Author contributions

RGG and SRM were involved in conceptualization and design.

SRM, RGG, MRA, CB, ABD, EM, EA, ST, RFS, MSB, AA, and RAL performed the experiments.

SRM, MRA, RFS, ABD, EM, EA, MSB, AA, RAL and ST provided experimental data and statistical analysis. SRM, MRA, RAL, ST and RGG wrote the manuscript.

SRM, MRA, CB, ABD, ST, RFS, LBP, MSB, AA, YS, RAL and RGG edited the manuscript.

